# Auditable AI agents for verifiable end-to-end scientific analysis

**DOI:** 10.64898/2026.11.25.769078

**Authors:** Yun Ma, Hangyu Zhou, Luyao Qin, Yuhua Liao, Jingqi Yang, Aiping Wu

## Abstract

Artificial intelligence (AI) agents are evolving from research assistants into end-to-end executors, yet their capacity to generate conclusions at scale outpaces rigorous verification. Here we introduce an auditability-centred paradigm for scientific automation, wherein trust relies not on generative capability or self-evaluation, but on whether claims can be independently traced to primary evidence, reviewed and corrected. Moving beyond post-hoc checks, we embed verification throughout the research lifecycle via four principles: grounded knowledge, bounded execution, evidence-backed results and independent verification. We instantiate this paradigm in EvoTrace for viral evolutionary analysis. Distilling peer-review critiques and methodological literature, we curated 13 principles, 236 pitfalls and 50 analytical units—validating 47 across a six-tier hierarchy—to encode methodological limits as executable constraints. Across simulated benchmarks, six viral datasets and an end-to-end Crimean–Congo haemorrhagic fever virus (CCHFV) study, claim–evidence auditing compiled evidence packages for all 233 analyses (flagging 26 incomplete evidentiary chains), while independent numerical and figure audits rectified 17 flaws, including three blocker-level errors. For CCHFV, EvoTrace intercepted a data-contradictory claim, recalibrating ‘no reassortment candidates’ to ‘no high-confidence candidates’. By co-delivering verification evidence, auditable automation shifts AI-driven science from autonomous generation to continuous scrutiny and correction.

## Introduction

Artificial intelligence (AI) is transitioning from an auxiliary research tool into an autonomous executor of the scientific process^1,2^. Beyond pattern recognition and generative prediction, AI systems now formulate hypotheses, design experiments, invoke tools and refine reasoning via feedback^2,3^. Whereas The AI Scientist automates the entire lifecycle from hypothesis generation to manuscript preparation^4^, Virtual Lab, AI co-scientist and Robin orchestrate multi-agent collaborative systems^5–7^. Similarly, Biomni integrates literature, databases and tools into a unified biomedical environment^8^, and Paper2Agent converts published papers and codebases into callable capabilities^9^. Scientific automation is thus shifting from isolated tasks to continuous, end-to-end workflows.

However, more complete research artefacts do not guarantee more reliable conclusions^10^: hallucinations, biases and irreproducibility persist even in integrated end-to-end workflows^2,10,11^. Systematic evaluations by ScienceAgentBench and PaperBench—assessing data-driven tasks and full-paper reproduction, respectively—reveal substantial agent limitations in real-world scenarios^12,13^. Recent events in mathematics have elevated this issue from capability benchmarking to a broader reckoning over scientific trust. In October 2026, OpenAI simultaneously released 722 AI-generated mathematical manuscripts, three of which were subsequently withdrawn owing to sign errors and other flaws^14,15^. Although verification standards vary across disciplines, this incident and the ensuing discourse expose a universal challenge: when AI mass-produces candidate findings, does the scientific community possess commensurate mechanisms for verification, comprehension and accountability^16^? Consequently, rather than chasing leaderboard gains, we address a more fundamental question: how should scientific trust be established within end-to-end autonomous research?

Analytical reliability depends on both the final output and the sequence of decisions leading to it^17,18^. Choices regarding data retention, outlier exclusion and method selection profoundly alter downstream analyses and interpretations^10^. Whereas traditional publications document static methods and conclusions, they rarely capture the dynamic decision trajectories of running agents. Furthermore, when planning, execution and self-checking share a context window, verification risks inheriting generative assumptions. Verification, therefore, cannot be deferred until report drafting; it must be shifted upstream to critical decision junctures and embedded directly into analytical execution.

This upstream shift requires decomposing complex research into discrete, verifiable units. Existing efforts have laid important groundwork: Paper2Agent converts published methods into tested, callable tools^9^; Scientific Agent Skills curates workflows into documented, scripted skill libraries to operationalize methodological knowledge^19^; and Claude’s engineering practices separate continuous-context execution from isolated-context independent judgement^20^. Yet callability and reusability do not equate to end-to-end trustworthiness. Analytical units require explicit preconditions and proof of correctness, and final claims must withstand scrutiny independent of generation.

This requirement points to a first principle: scientific trust cannot rest solely on the scale, autonomy or self-evaluation of a generative system, but must enable uninvolved parties to reconstruct evidence and test claims. Auditability is thus not a substitute for scientific truth, but a prerequisite for external scrutiny. Rather than serving as a post-hoc review appended to a final report, it must permeate four foundational layers: grounded knowledge, bounded execution, evidence-backed results and independent verification.

We translate this principle into two complementary mechanisms: execution-embedded verification and generation-independent auditing. From historical peer reviews, our system distils recurring failure modes, methodological boundaries and evidentiary standards to guide specialized agents; from methodological literature, it extracts expert-calibrated, composable skills with explicit validation tiers and applicability domains. During execution, these skills enforce quality constraints and log inputs, parameters and intermediate artefacts. Subsequently, a ‘claim-binding agent’ maps scientific assertions to these execution logs, verifying evidentiary chain integrity and flagging omissions or contradictions; critical flaws must be rectified or justified, then reviewed and signed off by a human expert. Finally, an isolated ‘auditing agent’ traverses the claim–evidence chain to recompute values, cross-check figures and track corrections. Verification thus spans low-level execution, claim–evidence anchoring and conclusion auditing, delivering both scientific conclusions and the complete evidentiary provenance required to scrutinize and correct them.

We evaluate this framework in viral evolutionary analysis—a domain characterized by heterogeneous inputs and tightly coupled stages, where subtle early deviations in quality control, phylogenetic rooting or temporal signal assessment can propagate downstream to distort tree topologies and evolutionary interpretations^21–23^. Simulated ground truths, curated reference builds and empirical datasets provide external benchmarks for quantitative evaluation^24–26^. Progressing from controlled benchmarks, we apply the full framework to an end-to-end study of Crimean–Congo haemorrhagic fever virus (CCHFV). Viral evolution serves as a concrete instantiation of our four principles rather than their boundary. Other disciplines can reconstruct domain-specific knowledge, skills and validation standards within the same four-tiered architecture, although cross-domain efficacy must ultimately be tested against field-specific ground truths, tasks and independent verification.

## Results

### An auditable architecture for end-to-end scientific analysis

As AI agents generate outputs at machine speed, manual evaluation cannot keep pace; yet generative capability does not equate to trustworthiness. Transitioning from AI-assisted research to end-to-end scientific analysis requires bridging the ‘trust gap’ between rapid production and reliable verification. We therefore propose an auditability-centred scientific automation framework that embeds the prerequisites for scientific trust directly into result production and verification (Fig. 1a). The framework follows four interlocking principles: grounded knowledge, ensuring traceable analytical rationales; bounded execution, confining analyses within defined applicability domains and operational constraints; evidence-backed results, establishing verifiable mappings from scientific claims to inputs, parameters, intermediate artefacts and final outputs; and independent verification, using agents uninvolved in generation to audit key findings and their evidentiary support. The first three principles establish the basis for inspection, whereas the fourth decouples verification from generation to prevent assumptions and errors from persisting through self-evaluation.

**Fig. 1.**
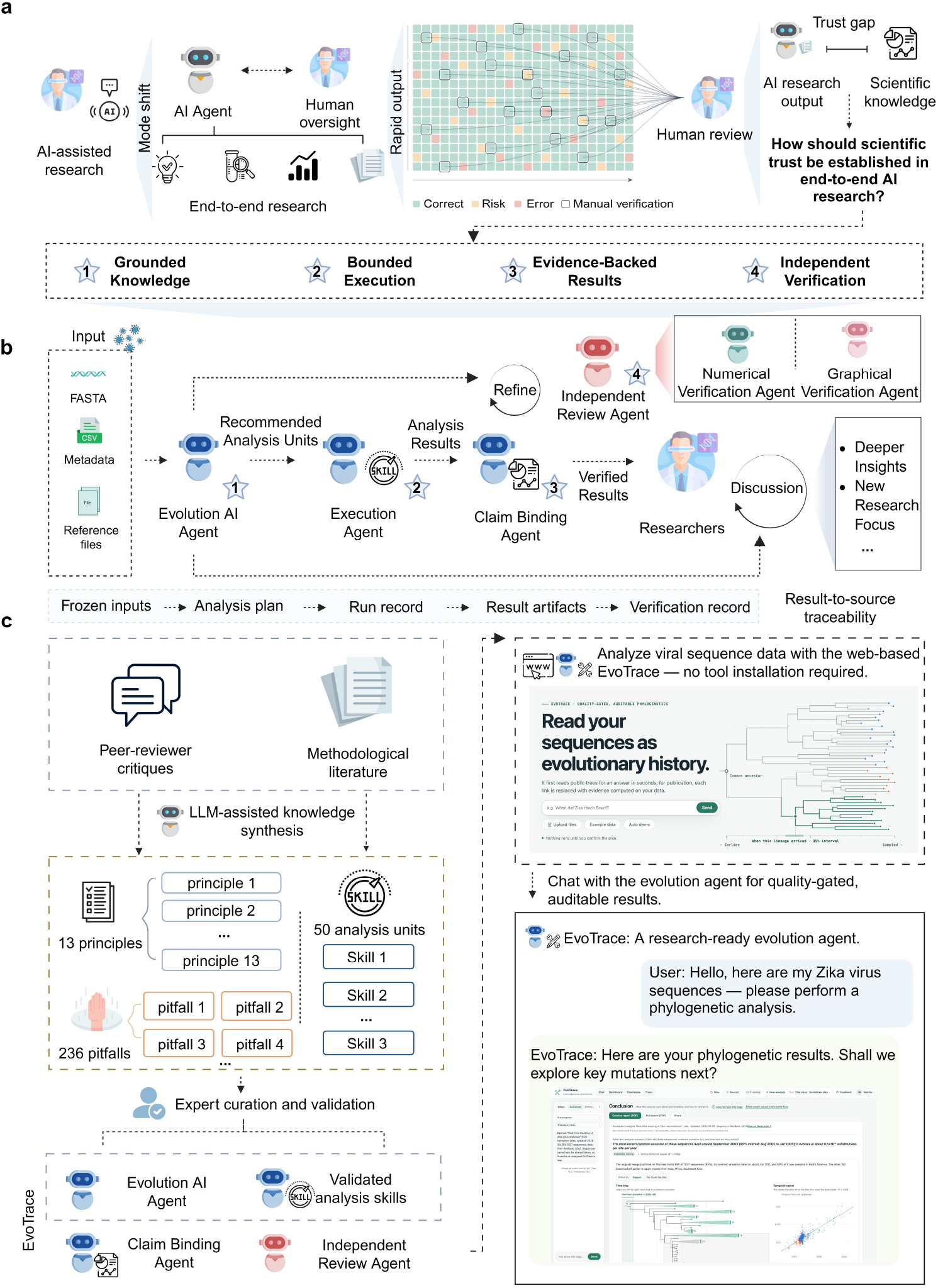
Auditable agent architecture for end-to-end scientific analysis. **a,** Trust gap in end-to-end AI research and four trustworthiness principles (★1–4 match core modules in b; coloured blocks are conceptual). **b,** Multi-agent workflow and verification dataflow: from task parsing (★1), controlled execution (★2), claim binding (★3) and dual-channel (numerical and graphical) blinded auditing with closed-loop correction (★4) to human–AI discussion, maintaining end-to-end provenance from results to raw data. **c,** Domain knowledge base construction (13 principles, 236 pitfalls and 50 analytical units), EvoTrace skill encapsulation and web interface (exemplified by Zika virus phylogenetics).

Four core workflow agents operationalize these principles, forming a complete evidentiary chain from final results back to source data (Fig. 1b). First, an evolutionary analysis agent parses research questions and inputs (such as sequences, metadata and reference files) to recommend analytical units. An execution agent then invokes corresponding skills to compute outputs under predefined constraints, and a claim-binding agent anchors research claims to specific supporting evidence. Subsequently, an independent verification agent coordinates with numerical and graphical verification agents for a blind audit: the former recomputes key metrics from underlying data, while the latter checks consistency across visual elements, annotations and analytical artefacts. Detected discrepancies trigger closed-loop refinement (‘Refine’) and re-inspection. By sequentially logging frozen inputs, analytical plans, execution logs, outputs and audit trails, the system achieves full result-to-source traceability. Researchers then interact with agents over these verified results and provenance records to derive deeper insights and new hypotheses. Independent verification thereby operates as an embedded control between generation and interpretation rather than a post-hoc review, strictly testing evidentiary support without equating audit passage to scientific truth.

Instantiating this framework for viral evolutionary research, we developed the domain system EvoTrace (Fig. 1c). By extracting recurring analytical flaws from peer reviews and task specifications from methodological literature—through large language model (LLM)-assisted knowledge synthesis combined with expert curation and validation—we built a domain library comprising 13 scientific principles (Supplementary Table 1), 236 analytical pitfalls (Supplementary Table 2) and 50 analytical units (Supplementary Table 3; 47 of which undergo tiered validation alongside 3 basic utilities). This knowledge is codified into workflow constraints, quality-control requirements and evidentiary standards empowering the four core agents. Deployed as an installation-free web platform (https://evotrace.ai), EvoTrace allows users to upload data and interact with agents to obtain quality-gated, auditable analyses. This implementation provides a unified testbed to sequentially evaluate knowledge-guided flaw detection, analytical unit reliability, end-to-end inference performance and the impact of independent verification on scientific conclusions.

### Peer-review critiques provide grounded knowledge for analytical judgement

Scientific analysis requires not only executing computations, but also judging whether conclusions withstand scrutiny. Whereas execution relies on explicit methodological rules, evaluation depends on tacit scientific judgement regarding when evidence fails to support a claim—weighing preconditions, alternative explanations and evidentiary sufficiency. Rarely codified in textbooks, such standards are repeatedly articulated in peer-review critiques. We therefore mined historical peer reviews to distil scattered critiques into high-level ‘judgement principles’ and specific ‘analytical pitfalls’, evaluating them across knowledge structure, temporal generalization and real-world failure coverage (Fig. 2).

**Fig. 2.**
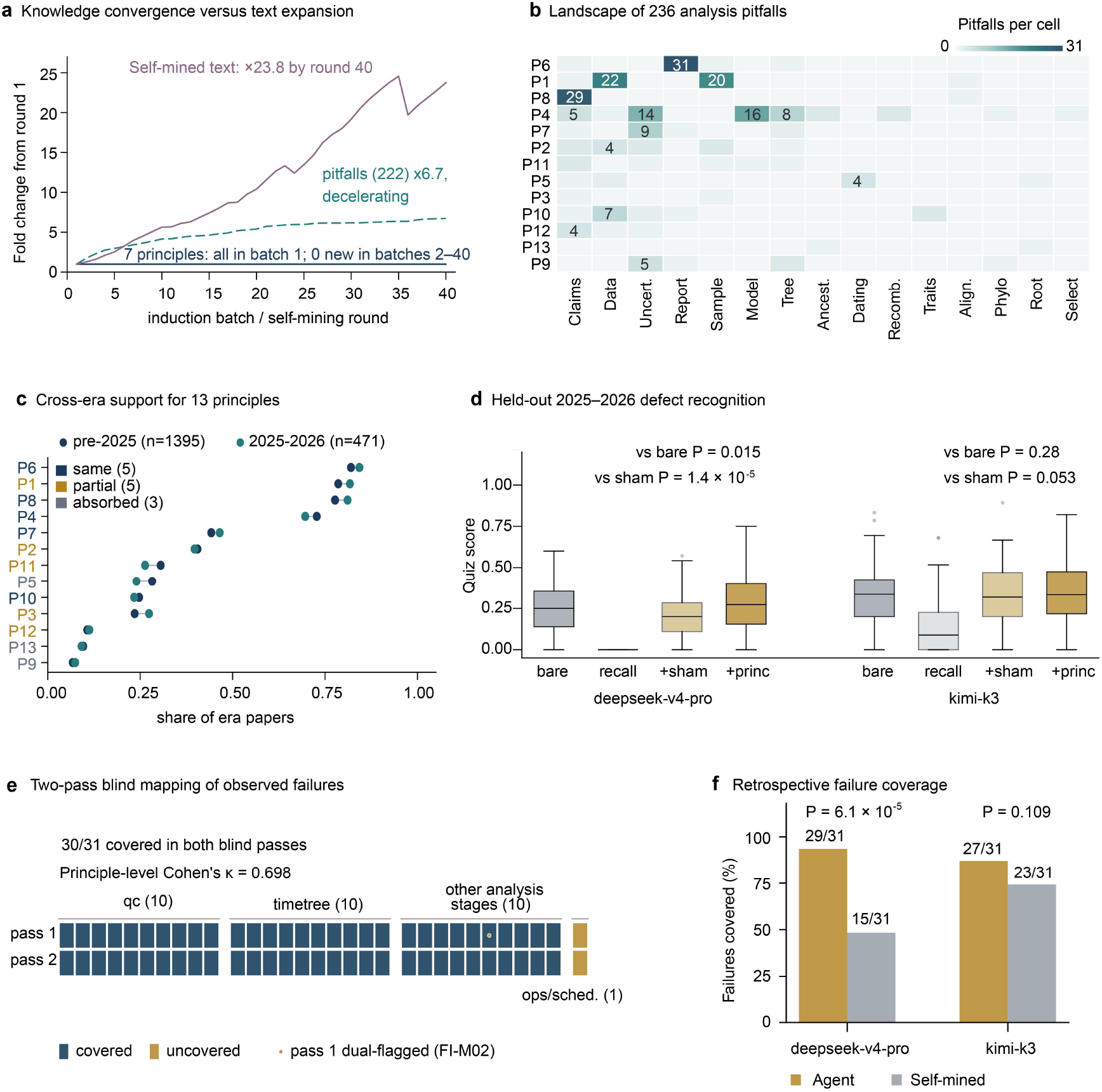
Grounded knowledge: architecture, cross-period support and defect-identification utility of review-induced knowledge. **a,** Normalized cumulative growth curves of batch-induced and self-mined knowledge. **b,** Heatmap of 236 pitfalls across 13 principles and 15 analytical stages. **c,** Paper support rates for the 13 principles across pre-2025 (n = 1,395) and 2025–2026 (n = 471) corpora and their mapping to earlier principles. **d,** Defect-identification scores under four prompting strategies (bare, Methods excerpt only; recall, metadata only; +sham, generic control; +princ, injected with seven earlier principles; n = 60 paired items per condition; boxes denote median and interquartile range (IQR), whiskers 1.5 × IQR and points outliers; one-sided paired Wilcoxon signed-rank test). **e,** Two-round blinded mapping matrix for 31 empirical failure units (principle-level Cohen’s κ = 0.698). **f,** Retrospective coverage of the 31 failure units by the review-induced (Agent) versus self-mined (Self-mined) checklists (one-sided exact McNemar’s test).

As the corpus grew, judgement principles rapidly converged and remained temporally stable, whereas specific pitfalls progressively refined. Across 40 incremental batches of pre-2025 corpora, all seven core principles (principles_2024) emerged in the first batch (Supplementary Table 4), while pitfalls increased from 33 to 222 before plateauing (Supplementary Table 5); conversely, unrefined self-mined baseline text expanded 23.8-fold by batch 40 (Fig. 2a). The full-period knowledge base comprises 13 principles and 236 pitfalls across 15 analytical stages, enriched in reproducibility, claim–evidence alignment and data quality (Fig. 2b). These 13 principles recurred consistently across pre-2025 (n = 1,395) and 2025–2026 (n = 471) corpora and aligned closely with the earlier seven principles (five direct matches, five partial matches and three subsumed; Fig. 2c). Moreover, substituting self-mined text with structured principles cut token overhead by 57.5%, achieving net cumulative savings after ∼1,328 calls including one-time construction costs (Extended Data Fig. 1).

In temporally split backtesting, early principles helped models detect methodological flaws in newer literature. Freezing principles_2024 against 60 held-out 2025–2026 test items, we conducted paired evaluations across unguided (bare), model self-recall (recall), generic control (sham) and principles_2024 conditions (Fig. 2d). Injecting principles_2024 significantly improved DeepSeek-V4-Pro over bare and sham controls, with median paired score gains of 0.0327 and 0.0775 (one-sided Wilcoxon signed-rank test, *P*= 0.015 and 1.4 × 10^−5^); corresponding Kimi-K3 gains were 0.0387 and 0.0243 (*P* = 0.278 and 0.053). Both models improved directionally, with statistical support concentrated in DeepSeek-V4-Pro.

Specific analytical pitfalls captured most real-world bare-model failures, outperforming self-mined knowledge. Recurring bare-model errors in real analyses consolidated into 31 failure units. Two-round blinded coding showed that curated pitfalls covered 30/31 units (principle-level Cohen’s κ = 0.698), missing only one low-level scheduling issue (Fig. 2e). Comparing peer-review-distilled against self-mined checklists, DeepSeek judged coverage at 29/31 versus 15/31 units (14 versus 0 discordant units; one-sided exact McNemar’s test, *P* = 6.1 × 10^−5^), and Kimi judged 27/31 versus 23/31 (5 versus 1 discordant units; *P* = 0.109; Fig. 2f). Peer-review-derived knowledge therefore combines temporal stability, low token overhead and broad failure coverage; we next test its protective value during live end-to-end execution.

### Composable analytical units define verifiable execution boundaries

Because data filtering, method selection and parameterization are interdependent in complex analyses, local biases can propagate along pipelines, obscuring their origins in final reports. We therefore decompose workflows into composable analytical units, shifting verification upstream to key generative steps: each unit performs an independently inspectable task while logging preconditions, validation results and open issues; once assembled, cross-step event recovery and diagnostics are evaluated. Agents accordingly invoke methods that are explicitly bound to validation evidence and operational limits. In EvoTrace, we built 50 composable units across 13 evolutionary domains and established a six-tier validation hierarchy across all 47 non-utility units (spanning unit tests, diagnostics, cross-method comparisons, planted defects, simulated ground truths and official reference builds, with highest tiers covering 22, 1, 1, 2, 17 and 4 units, respectively); the 137 validation records comprise 134 passes, 2 below preregistered thresholds and 1 robustness report (Fig. 3a).

**Fig. 3.**
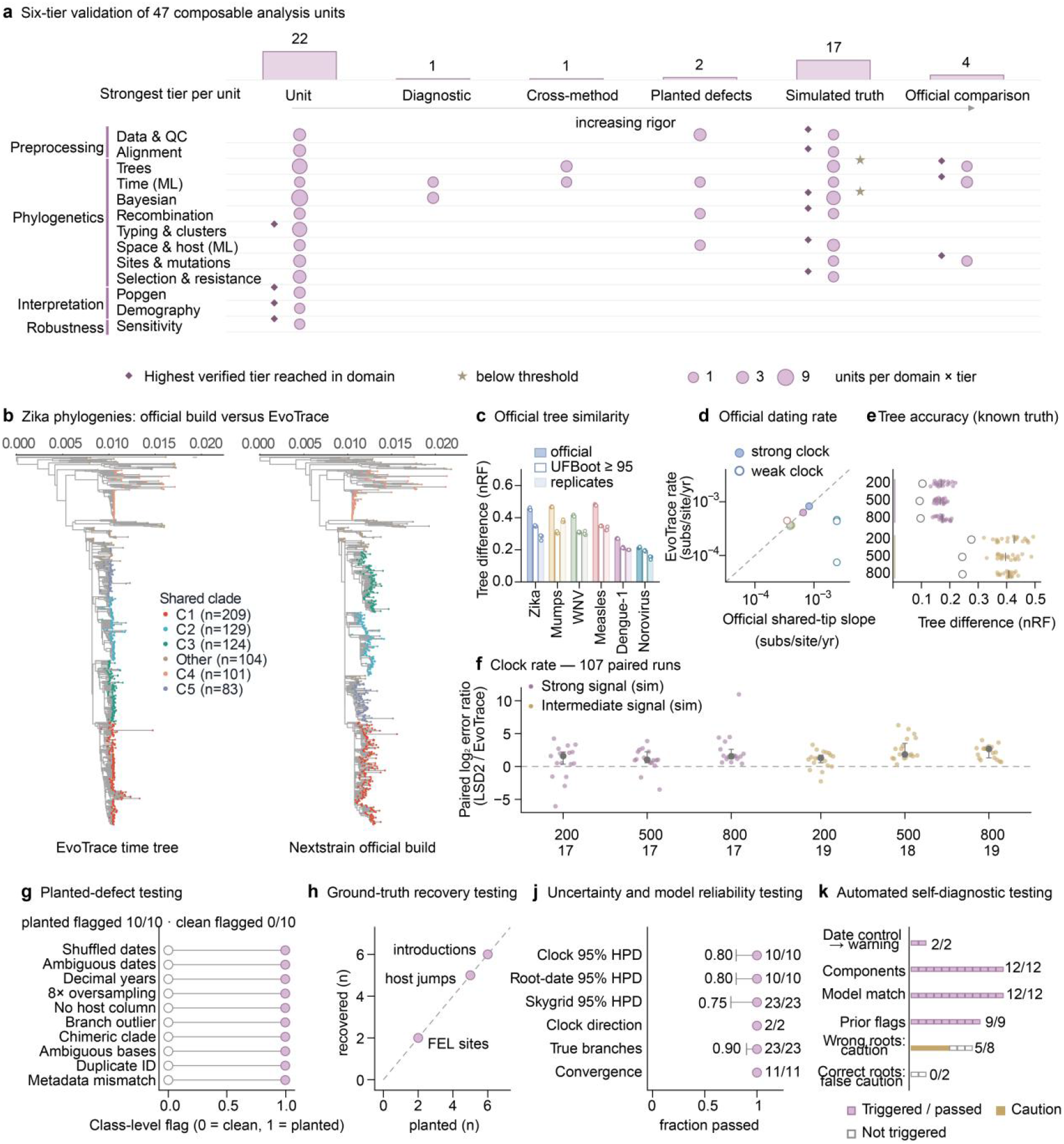
Bounded execution: hierarchical validation of composable analytical units. **a,** Highest validation tiers for 47 non-foundational units across 13 domains (137 evidence records: 134 passed, 2 sub-threshold [asterisks], 1 robustness report; diamonds denote domain maxima). **b,** Topological comparison of the EvoTrace Zika virus time tree against the official Nextstrain reference (750 shared tips). **c,** Normalized Robinson–Foulds (nRF) distances between EvoTrace and reference trees across six empirical viral datasets (mean ± s.d., n = 3 independent runs). **d,** EvoTrace clock rates versus reference root-to-tip slopes (n = 3 independent runs; solid and open symbols, strong and weak temporal signals; dashed line, y = x). **e,** nRF distances to ground truth across 120 simulated trees (purple and yellow dots, individual runs with strong and moderate temporal signals; vertical ticks and open circles, group medians before and after collapsing UFBoot < 95% branches; 0 denotes identity). **f,** log_2_ relative clock-rate error ratios of LSD2 to EvoTrace in simulations (positive values favour EvoTrace; median and 95% CI; left to right, n = 17, 17, 17, 19, 18 and 19 valid paired runs). **g,** Category-level detection across ten injected defect classes (10/10) versus clean controls (0/10 false alarms). **h,** Cross-step recovery of simulated events and selected sites (six introductions, five host jumps, two FEL sites; dashed line, y = x). **j,** Ground-truth coverage and convergence pass counts across six Bayesian benchmarks. **k,** Trigger frequencies for model-assumption diagnostics (date shuffling, 2/2; model misspecification and compositional anomalies, 12/12 each; prior-dominated parameters, 9/9; misrooting, 5/8; correctly rooted controls, 0/2 false alarms; label i omitted).

Unit-level verification ensured accurate foundational inference and traceable upstream biases. For Zika virus (750 shared tips), EvoTrace’s time-scaled tree closely matched the official Nextstrain reference, recovering the C1–C5 clades (Fig. 3b). Across six Nextstrain viral datasets, topology converged stably, with mean normalized Robinson–Foulds (nRF) distances of 0.215–0.479 to official trees (0.152–0.379 between replicates; Fig. 3c). Under strong temporal signal, clock rates differed from official slopes by −12.48% to +0.51% (0.51% for Zika), while measles and norovirus were accurately flagged as weak-signal boundary cases (Fig. 3d). Unit logs also isolated preprocessing impacts: masking untranslated region noise in West Nile virus raised patristic distance correlation (Pearson’s r) with the official tree from 0.402 to 0.994 (Extended Data Fig. 2a–c, e, f). In simulated benchmarks, topological accuracy tracked signal strength (median nRF of 0.166–0.174 and 0.396–0.427 for strong and moderate signals; Fig. 3e and Extended Data Fig. 3h), with 118/120 clock rates meeting criteria. Across 107 paired comparisons with LSD2, LSD2’s median clock rate errors were 2.04–6.34-fold higher than EvoTrace’s (all six 95% confidence intervals > 1; Fig. 3f), and EvoTrace produced more accurate time to the most recent common ancestor (tMRCA) estimates in 60% (64/107) of runs (Extended Data Fig. 3a, e).

Assembled into multi-step pipelines, cross-step verification blocked silent error propagation, safeguarding complex event recovery and assumption checks. Across 10 classes of planted pipeline defects, detection was 100% with zero false positives in clean controls (Fig. 3g). In coupled evolutionary scenarios, the pipeline recovered all simulated cross-regional introductions (6/6), host jumps (5/5) and positively selected sites (2/2) without false positives (Fig. 3h). Beyond passing all six Bayesian ground-truth checks—95% highest posterior density (HPD) coverage, Skygrid dynamics, model selection and chain convergence (Fig. 3j)—diagnostic units captured date shuffling (2/2), model misspecification and compositional heterogeneity (12/12 each), prior domination (9/9) and erroneous rooting (5/8; 0/2 false alarms on correct roots; Fig. 3k), extending verification to statistical premises.

Binding preconditions and gating rules within units enabled agents to expose failures rather than force outputs on out-of-bounds data. Upfront quality control and dating triage intercepted all high-N sites (24/24), duplicates (48/48) and misgroupings (9/9), alongside 53/120 (44.2%) date errors (Extended Data Fig. 3f) and all 30 signal-free datasets (0/150 misclassifications; Extended Data Fig. 3b), confining inference to valid regimes (82.2% coverage of 90% tMRCA HPD in valid runs; Extended Data Fig. 3c). Plausibility gates captured 12.1% (26/215) of aborted simulated dating runs (Extended Data Fig. 3g) and exposed empirical norovirus dating divergence (to year −3653.16; Extended Data Fig. 2d), while runtime scaled predictably with sample size (log–log slope = 1.86; Extended Data Fig. 3d). Together, composable units anchor successes, failures and uncertainties to specific steps, establishing traceable execution boundaries.

### Evidence binding links analytical claims to verifiable evidence

To bridge computation and reporting, evidence binding links analytical claims to underlying data, intermediate steps and output files, withholding delivery when evidentiary chains are incomplete or unverified and strictly separating ‘results generated’ from ‘evidence sufficient for delivery’ (Fig. 4a, b). Our Claim Binding Agent integrates raw inputs, analytical steps, stage-gate quality controls, outputs and execution logs into a provenance record, sequentially performing rule checks, agent self-inspection and read-only retrieval of independent recalculations. Blocking issues require resolution or justification before re-inspection, and final delivery mandates human sign-off, producing an evidence bundle with reported claims, verification records and delivery status for each analysis (Fig. 4a, b).

**Fig. 4.**
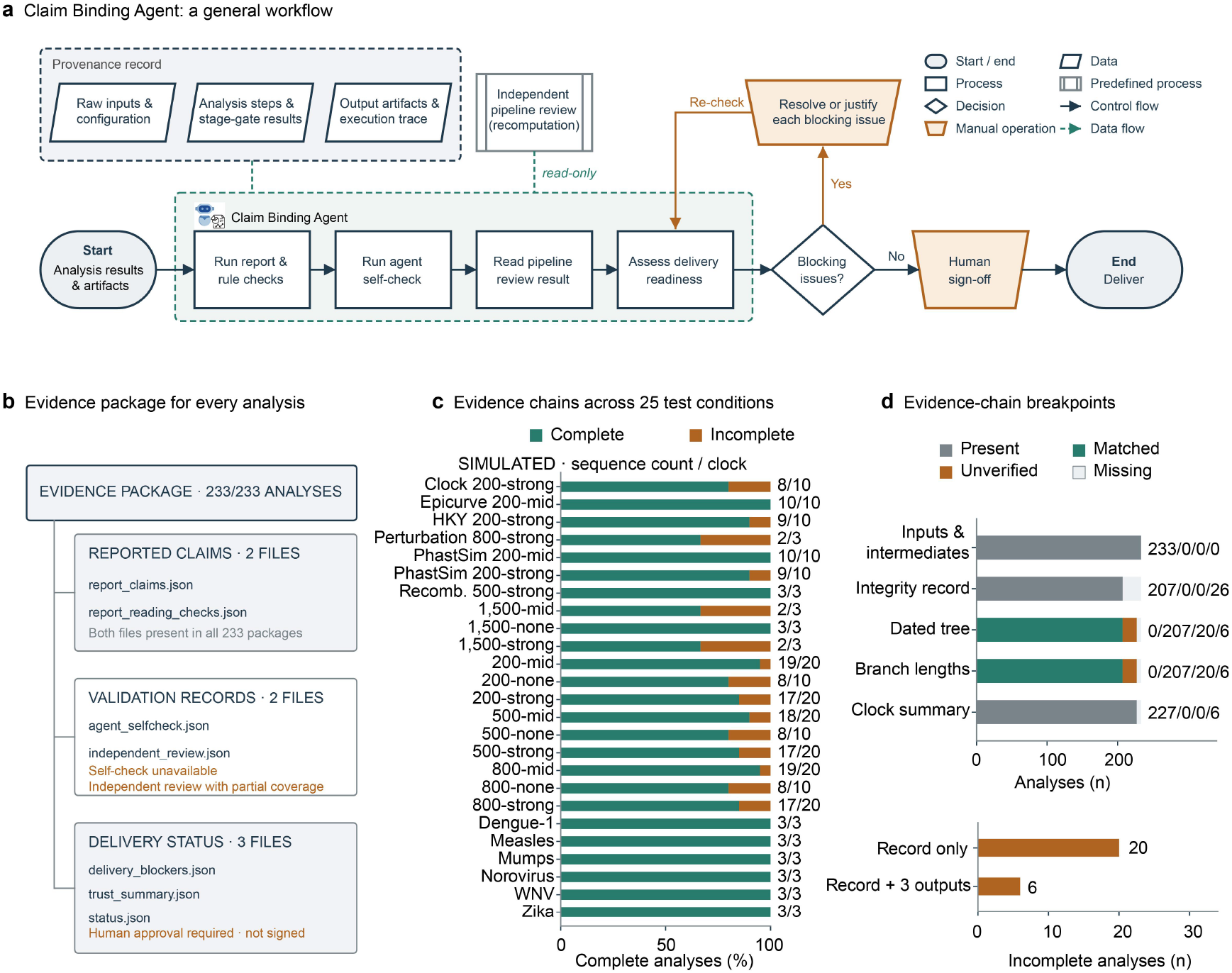
Evidence-bound results: evidence-chain binding and integrity tracking of analytical outputs. **a,** Evidence verification, anomaly blocking and human sign-off by the claim-binding agent. **b,** Evidence-bundle structure and overall verification status across 233 analytical tasks. **c,** Evidence-chain completeness rates across 25 test conditions (215 simulated and 18 empirical runs). **d,** Breakpoint types and causes for the 26 incomplete chains (from c): 20 generated dated time trees, branch lengths and clock summaries but lacked integrity records for tamper checks; 6 lacked both records and outputs (green, orange, grey and white denote checksum-matched, present but unverified, present only and missing, respectively, separating ‘output generated’ from ‘output verifiable’).

Across 233 analyses spanning 25 test conditions (215 simulated, 18 empirical), 207 met run-level integrity rules (resolvable inputs, complete files and matching hashes for time-scaled trees and branch lengths), while 26 were retained as incomplete evidence chains (Fig. 4c). Of 1,290 reported claims, 1,212 had complete evidence chains; of 466 output links, 414 passed hash verification and 52 lacked reference hashes, with zero mismatches (Fig. 4d). All input and intermediate files remained intact; of the 26 incomplete analyses, 20 lacked only integrity records and 6 lacked time-scaled trees, branch lengths and clock summaries (Fig. 4d), pinpointing missingness to exact files. With agent self-inspection disabled and independent replication only partially covered, all 233 analyses were held unsigned pending human approval (Fig. 4b). Because current chains reflect run-level file associations rather than field-level mappings, chain completeness indicates traceable file dependencies rather than completed independent verification or biological correctness.

Using the 18 empirical analyses, two controlled blinded experiments evaluated 234 paired replicates (144 single-defect, 90 cumulative-defect; 216 unique condition replicates after merging shared controls; Extended Data Fig. 4a). Detection and exact localization reached 100% for both single- and cumulative-defect replicates (126/126 and 72/72), with all controls passing (18/18; Extended Data Fig. 4b). Claim screening retained all 103 unaffected claims and intercepted all 721 and 412 affected claims across the two arms (100% accuracy: 824/824 and 515/515; Extended Data Fig. 4c). From cumulative defect levels 0 to 4, anomalous (missing or unverified) core file proportions rose linearly (0, 0.125, 0.250, 0.375 and 0.500; Extended Data Fig. 4d). Evidence binding therefore reliably intercepts predefined structural defects and blocks compromised claims from delivery, while leaving semantic logic flaws and domain-level misinterpretations to downstream review.

### Independent auditing enables defect localization, boundary quantification and delivery remediation

To prevent local error propagation across multi-step workflows and avoid assumption inheritance from same-context self-checks, we decoupled auditing into independent numerical and figure verification channels (Fig. 5a). An orchestrator agent freezes the audit baseline and assigns blinded tasks: a numerical agent rebuilds summary tables from raw artefacts to check formulas and definitions, while a figure agent reverse-maps graphical elements to underlying data. Both channels submit evidence independently; the orchestrator adjudicates severity against raw records and drives traceable, closed-loop remediation via a versioned ledger.

**Fig. 5.**
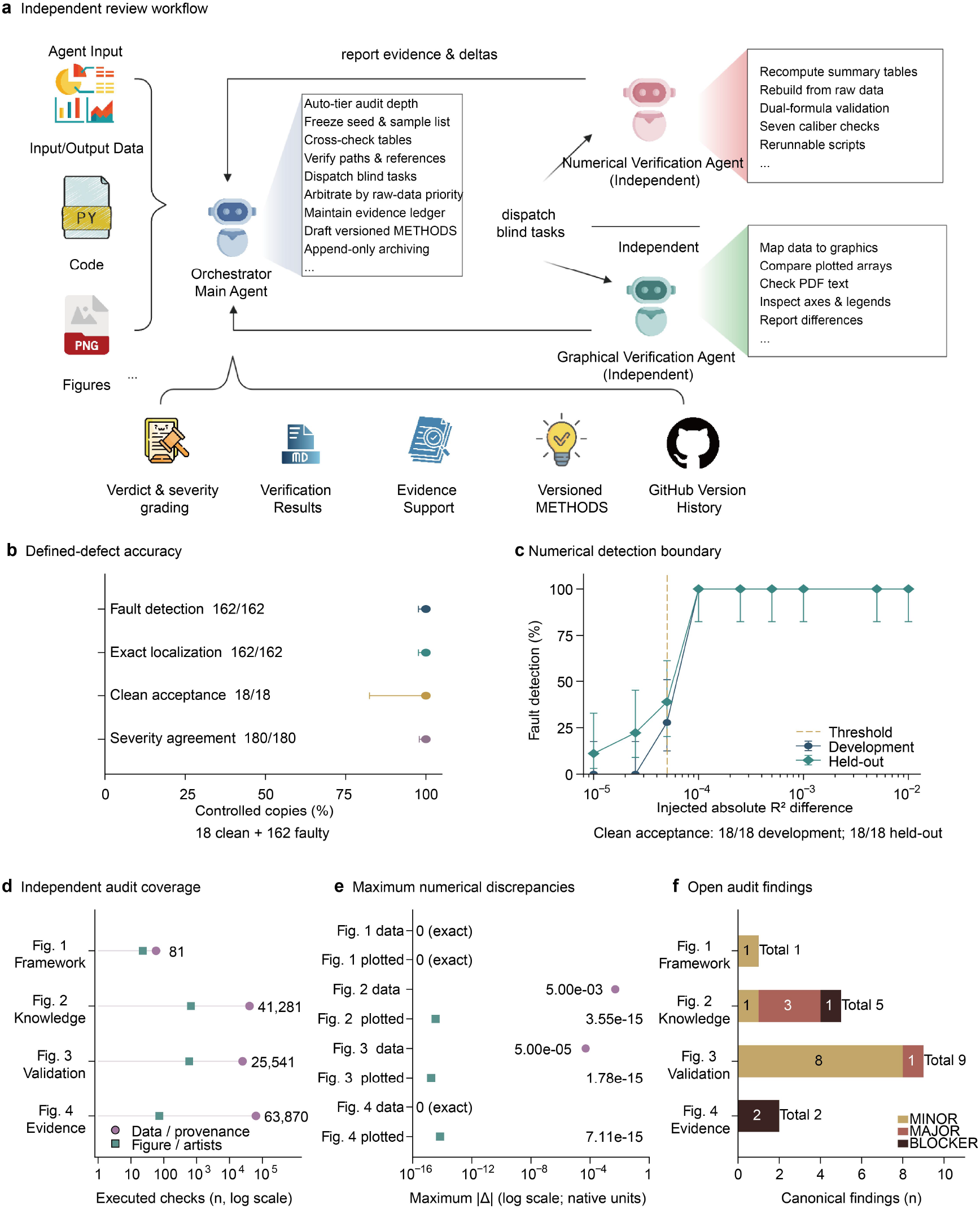
Independent auditing: architecture, benchmark performance and initial audit results of the auditing workflow. **a,** Independent auditing workflow: the orchestrator freezes baseline artefacts, assigns blinded tasks to decoupled numerical and graphical agents, arbitrates dual-channel evidence against raw records, and outputs severity ratings, an evidence ledger and versioned logs. **b,** Detection accuracy in controlled defect injections (n = 180 replicas from 18 empirical runs across six viruses: 18 clean controls and 162 single-defect replicas across nine classes). Dots and error bars show accuracy (all 100%) and two-sided 95% Wilson CIs. **c,** Numerical detection boundary across nine absolute R^2^ drift tiers (10^−5^ to 10^−2^) on 36 four-decimal R^2^ chains (n = 360 trials: 324 injections, 36 clean controls). The threshold (5.02×10^−5^, vertical dashed line) was set on the development split (dengue, measles and mumps; n = 18 chains per tier) and tested on the hold-out split (norovirus, WNV and Zika; n = 18 chains per tier; clean-control acceptance, 18/18 in both). Points and error bars show detection rates and two-sided 95% Wilson CIs. **d,** Initial audit check coverage (log scale) across four evidence lines: Fig. 1 framework (n = 81), Fig. 2 knowledge and Extended Data Fig. 1 (n = 41,281), Fig. 3 validation and Extended Data Figs. 2 and 3 (n = 25,541), and Fig. 4 evidence (n = 63,870), totalling 130,773 checks across data/provenance (purple circles) and figure/element (cyan squares) channels. **e,** Maximum absolute deviations (∣Δ∣, log scale in native units) across 39,099 direct numerical comparisons; all passed within pre-set tolerances, with non-zero values reflecting two- or four-decimal rounding bounds or floating-point limits. **f,** Initial audit open findings (n = 17), stratified by evidence line and severity (10 MINOR, 4 MAJOR, 3 BLOCKER).

In blinded defect tests across six viruses (18 cases), the workflow detected and localized all 162 defective replicates across nine error classes (100%; two-sided 95% CI: 97.7–100%), cleared all 18 clean controls (100%; 82.4–100%) and matched preset severity ratings across all 180 samples (100%; 97.9–100%; Fig. 5b). To bound numerical sensitivity, we injected nine absolute-drift levels (10^−5^ to 10^−2^; 360 trials) into 36 four-decimal R^2^ chains. Using a 5.02×10^−5^ threshold calibrated on three viruses, evaluation on three held-out viruses showed partial detection at 10^−5^ to 5×10^−5^ due to four-decimal rounding residuals (5/54 development, 13/54 held-out), but 100% detection at ≥10^−4^ in both sets (18/18 per tier; held-out overall 74.7%, 121/162) with zero false positives in controls (Fig. 5c), separating true numerical errors from rounding noise.

In a delivery audit of our own Figs. 1–4 and Extended Data Figs. 1–3, an initial 130,773-check pass (Fig. 5d) included 39,099 matched-definition numerical comparisons, all passing within tolerance; maximum absolute deviations (0 to 5.00×10^−3^) strictly matched decimal-rounding bounds or floating-point limits (Fig. 5e). The initial audit also caught 17 structural and provenance defects across four evidence lines (10 MINOR, 4 MAJOR, 3 BLOCKER; 1, 5, 9 and 2 per line), involving label-definition misalignments, missing replication evidence and hash-manifest mismatches (Fig. 5f). After targeted fixes and re-auditing, checks rose to 131,427 (Extended Data Fig. 5a), precision across eight numerical chains held steady (Extended Data Fig. 5b) and open defects dropped to zero (Extended Data Fig. 5c). Notably, this framework excludes free-text semantics, pixel-level rendering and biological assumption review; closed-loop auditing ensures rigorous computational provenance rather than biological truth.

### EvoTrace improves analytical reproducibility, efficiency and auditability under identical foundation models and compute budgets

To isolate framework contributions from backbone capability, we benchmarked EvoTrace against the generalist agent Biomni and unconstrained Kimi-K3 under an identical Kimi-K3 backbone and compute budget (32 threads, 24 h limit) across six Nextstrain viral datasets and three random seeds (*n* = 18 per group; Fig. 6a).

**Fig. 6.**
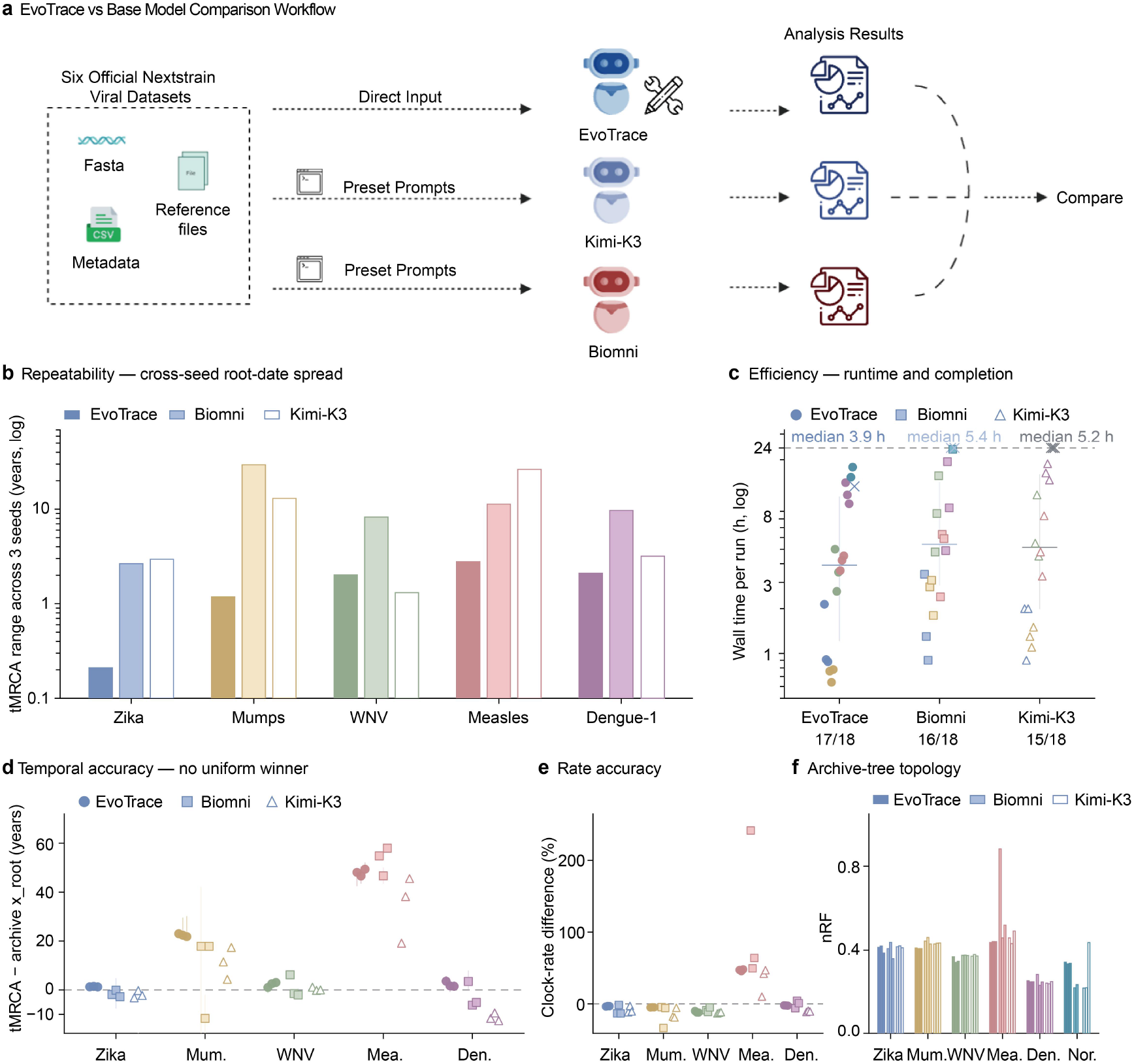
Three-way benchmark of EvoTrace, Biomni and unguided Kimi-K3. **a,** Experimental design (6 viral datasets × 3 methods × 3 seeds, n = 18 per group; unified Kimi-K3 backbone, 32 threads, 24-h limit; independent second-round EvoTrace runs; Nextstrain archived trees and rates as consistency references). **b,** Cross-seed (n = 3) tMRCA ranges across the five primary datasets. **c,** Runtime and completion counts across all 54 runs (dots, completed runs; horizontal and vertical lines, median and IQR; crosses, control time-outs or EvoTrace quality-gate aborts). **d,** tMRCA point-estimate deviations and compliant TreeTime 90% HPD intervals across the five primary datasets (45 runs; five Biomni runs show point estimates only due to missing or non-compliant intervals; dashed line, zero deviation). **e,** Clock-rate percentage deviations from archived rates (n = 45). **f,** Unrooted nRF distances on shared tips (n = 53; one Biomni norovirus run produced no tree; norovirus shown as a boundary case and excluded from testing). Panels b and c use log_10_ y-axes. Completion rates: two-sided Fisher’s exact test (n = 18 per group); ranges (b, n = 5) and primary-dataset runtimes (c), absolute deviations (d) and nRF distances (f; all n = 15 pairs): two-sided exact paired Wilcoxon signed-rank tests (unadjusted *P*values).

EvoTrace most notably reduced run-to-run variability. Across five datasets yielding comparable dating estimates, EvoTrace kept cross-seed tMRCA ranges to 0.21–2.78 years, outperforming Biomni on all five (2.65–29.51 years) and unconstrained Kimi-K3 on four (1.31–26.39 years; Fig. 6b), while maintaining highly consistent clock rates (three-run max/min ratio: 1.004–1.025). Structured domain protocols thereby constrain stochastic variance in the foundation model’s exploration trajectories.

EvoTrace also improved completion rates, efficiency and proactive quality control, completing 17/18 runs (versus 16/18 and 15/18) with a 3.91 h median runtime—28% faster than Biomni (5.40 h; paired Wilcoxon *P*= 0.055) and 24% faster than unconstrained Kimi-K3 (5.15 h; *P*= 0.0026) (Fig. 6c). Whereas every incomplete baseline run exhausted the 24-h wall-clock budget, EvoTrace’s sole unfinished norovirus run was deliberately halted by a built-in quality gate upon detecting a non-positive clock rate.

Overall, EvoTrace’s evolutionary estimates better matched archived Nextstrain references. While no method dominated all dating and rate metrics (Fig. 6d, e), EvoTrace yielded the lowest overall median absolute tMRCA deviation across 15 paired runs (3.04 versus 6.18 and 9.39 years; *P*> 0.2) and significantly lower normalized Robinson–Foulds (nRF) topological distances in 11/15 pairs versus Biomni (*P*= 0.008) and 12/15 versus unconstrained Kimi-K3 (*P*= 0.015) (Fig. 6f).

Beyond numerical metrics, EvoTrace excelled in controlled delivery and auditability. All 17 completed reports accurately extracted run-specific dating parameters and flagged weak temporal signals, whereas baselines frequently misreported internal nodes as roots, spliced intervals across runs, mislabelled confidence levels or concealed massive sequence pruning. While automated checks still require independent review (for example, an erroneous genogroup exclusion in norovirus), this comparison confirms that EvoTrace delivers faster, highly reproducible, reference-aligned and fully auditable results.

### An auditable CCHFV analysis loop links evolutionary hypotheses to evidence

Finally, using Crimean–Congo haemorrhagic fever virus (CCHFV), we illustrate how EvoTrace advances baseline phylogenetics into testable scientific hypotheses (Extended Data Fig. 6a, b). Following S-, M- and L-segment phylogenetic reconstruction, the agent autonomously generated and tested three progressive hypotheses. First, cross-segment screening detected 87 reassortment candidates among 377 resolvable multi-segment isolates (23.1%, conservative lower bound), with all 14 high-confidence candidates confined to Asia and Africa (Extended Data Fig. 6c, d). Second, European introduction reconstructions strongly supported a single eastern introduction (Russia to Turkey) across all three segments (80–100% bootstrap). Conversely, by the modal introduction count across 100 UFBoot replicates, multiple western introductions (Senegal to Spain/France) appeared only in the L segment, while other segments diverged in frequency and origin, reflecting small-sample limitations (Extended Data Fig. 6e). Third, isolate-level permutation tests revealed significant lineage–vector preferences—Europe 2 enriched in Rhipicephalus (7/8, q = 0.0023) and Europe 1 (S segment) in Hyalomma (19/25, q = 0.0437)—while explicitly bounding sampling confounders (Extended Data Fig. 6f). The full workflow passed independent auditing (Extended Data Fig. 7), achieving zero deviation across 133 ledger checks and two numerical recalculations. Auditing also blocked a BLOCKER phrasing flaw that omitted Turkish low-confidence candidates, enforcing revision to ‘no high-confidence candidates’ before release. In doing so, EvoTrace unifies hypothesis generation, quantitative testing and traceable error correction within one trustworthy framework.

## Discussion

As artificial intelligence begins generating ostensibly complete scientific reports, how can autonomous science establish trust? Here we show that trust cannot rest solely on foundation-model capability or superficial report completeness, but must be anchored in verification mechanisms embedded throughout the research process. To address this challenge, we structured an agentic workflow around four tenets: provenance-grounded knowledge, bounded execution, evidence-backed claims and independent verification. Mirroring core scientific norms, this ensures that claims allow independent reconstruction of evidence, validity testing and bias correction. Auditability does not guarantee correctness, but provides the prerequisite for scrutinizing conclusions and systematically correcting errors. Validating this strategy in molecular evolution—a field with long inferential chains and high uncertainty—EvoTrace embeds these principles across sequence quality control, phylogenetics and hypothesis testing, transforming black-box generation into an inspectable, revisable workflow.

Instilling domain norms into autonomous systems first requires expert priors. Historical peer-review corpora encode both explicit methodological rules and implicit judgements on applicability premises, alternative explanations and evidentiary sufficiency—guiding both how to run analyses and when results cannot sustain a conclusion. Yet distilling experience into operational rules does not fully encode complex scientific judgement. While retrospective tests show that these principles capture common past failures, they cannot guarantee interception of novel biases. Moreover, temporal-split evaluations reveal that performance gains from identical domain rules vary substantially across foundation models, and rubric scoring by a single model family may carry stylistic preferences; distilled rules therefore require cross-model or human-calibrated re-validation as foundation models evolve.

Given these limitations, our stratified evaluation decouples task completion, execution stability and inferential reliability. Unit verification isolates execution faults, whereas simulated and empirical benchmarks expose vulnerabilities in uncertainty estimation and weak temporal signals. Crucially, evaluation revealed a key blind spot: while protocol constraints markedly improve topological consistency across replicate runs, execution consistency does not equal inferential accuracy. Workflow stability can mask systematic bias. Reliable scientific automation thus requires both stable execution and an ability to recognize the applicability boundaries of data and methods.

To operationalize this boundary awareness, we placed a verification gate between result generation and conclusion delivery. Evidence binding packages claims with raw inputs, intermediate quality metrics and run logs into traceable bundles, withholding delivery if evidence is missing or quality gates fail, while retaining human sign-off for final release. To mitigate multi-step error propagation and ‘hypothesis inheritance’ during same-context self-checks, we decoupled auditing into independent, double-blind numerical and visual channels. These channels independently recompute statistics from raw artefacts and reverse-map visual elements to source arrays against locked baselines, triggering closed-loop remediation. In our CCHFV analysis, auditing caught preliminary over-inference and forced a substantive narrowing of final claims rather than adding superficial caveats. Still, architectural decoupling cannot fully erase shared foundation-model blind spots, nor can multi-agent consensus replace external evidence. Verification aims not to pass every hypothesis, but to calibrate claim strength to evidentiary weight; narrowing or retracting under-supported claims embeds scientific self-correction into automation.

Beyond molecular evolution, this architecture provides a template for other long-chain inferential sciences by reconstructing domain priors, bounding analyses, binding evidence and enforcing independent audits. However, just as model upgrades require re-testing, architectural transferability does not guarantee unconditional efficacy; each discipline must validate the approach against domain-specific benchmarks and compute budgets. Ultimately, autonomous scientific systems must deliver not only conclusions, but the complete evidentiary chain needed for others to test, question and revise them.

## Methods

### Peer-review corpus and knowledge induction

Applying predefined deduplication, cleaning and segmentation criteria to 4,753 candidate papers across 13 journals and open-review platforms, we selected 2,302 openly reviewed articles (1,395 with valid critiques from ≤ 2024; 471 from 2025–2026), yielding 29,347 valid critique units. Initial induction on 287 articles from eLife, F1000Research and Wellcome Open Research—combining model-assisted extraction with expert curation—produced a knowledge base of 15 analytical stages, 13 scientific principles and 236 analytical pitfalls. Another 47 analytical units with executable verification evidence were extracted from the Methods sections of 839 articles into the final ledger.

Temporal re-induction used only the ≤ 2024 corpus (21,511 in-scope critique units). With global labels masked, the model ingested 1,600 critique units from 93 articles in batches under a fixed random seed (20261020), stopping after ≥ 40 batches and 15 consecutive batches without new principles. This yielded 7 early principles and 222 pitfalls, which were used to classify and map the remaining early corpus (as consistent, partially corresponding or absorbed/merged). An autonomous mining baseline conducted 40 cumulative mining rounds on the same corpus.

### Temporal holdout examination and historical failure evaluation

The temporal holdout test comprised 60 questions and 665 ground-truth points derived from 60 articles (2025–2026). DeepSeek-V4-Pro (temperature = 0) and Kimi-K3 (temperature = 1) answered under four conditions: bare (question plus Methods excerpts), recall (bibliographic metadata only), sham (generic control text) and +principles (the 7 frozen early principles).

Anonymized responses were blind-scored by two independent DeepSeek-V4-Pro instances (temperature = 0). Ground-truth points scored 1 (hit), 0.5 (partial hit) or 0 (miss), while semantically independent irrelevant or invalid critiques counted as false alarms (FA). Per-question scores were computed as:

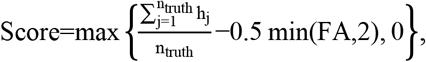

where n_truth_ is the total ground-truth points and h_j_ is the hit score for point j. Discrepancies ≥ 0.5 between raters triggered adjudication by a third instance; otherwise, scores were averaged. Paired per-question differences between +principles and bare or sham were evaluated using one-sided Wilcoxon signed-rank tests.

Thirty-one historical failure units were mapped to existing pitfalls via two rounds of independent blind coding to compute principle-level Cohen’s κ. Retrospective semantic coverage across the two knowledge types was judged by DeepSeek and Kimi and compared via one-sided exact McNemar’s tests. Using API-logged token usage and downstream character heuristics (1 token per CJK character; 0.25 tokens per non-CJK character), we derived the cost break-even point assuming fixed per-run cost and linear accumulation.

### EvoTrace architecture and stratified verification

EvoTrace links task parsing, bounded execution, evidence binding and independent auditing into a unified evidence chain. It executes protocol-driven quality control, deduplication, alignment, phylogenetic inference and reporting, saving input snapshots, parameters, logs and intermediate artefacts per run while binding inputs, artefacts, summary tables and final figures via SHA-256 hashes. In an isolated context, the native auditor recomputes metrics from raw artefacts and checks artefact–claim consistency (all reports require human sign-off).

Spanning 13 subdomains, the 47 analytical units are stratified by verification externality into six evidentiary tiers (unit tests, diagnostics, cross-method comparisons, planted defects, simulated ground truth and official reference validation). Pipeline-level tests cover planted defects, introductions, host jumps, fixed-effects likelihood (FEL) site recovery^27^ and Bayesian ground-truth checks. BEAST X (v.10.5.0)^28^ tests evaluated clock rate, root date and Skygrid recovery^29^ via 95% highest posterior density (HPD) intervals, requiring ESS ≥ 200, R^≤ 1.01 and tree-set ASDSF ≤ 0.01 for convergence. Structured coalescent validation ran dual chains in BEAST 2 (v.2.7.7)^30^ with MASCOT (v.3.0.7)^31^ (10% burn-in, stopping early upon convergence). Self-diagnostics included date shuffling, model-component checks, prior-dominance alerts and root-placement warnings.

### Simulation and Nextstrain benchmarking

Simulated ground-truth trees were generated via a birth–death sampling process (λ = 1.25, ψ = 1.0, R_0_ = 1.25), evolving 10-kb sequences under GTR+F+Γ4 in AliSim^32^. The primary grid (150 runs) crossed 200, 500 and 800 tips with strong (8×10^−4^ substitutions per site per year, 20 replicates), moderate (2×10^−4^, 20 replicates) and null temporal signals (permuted tip dates, 10 replicates). Hardening tests (65 runs) covered the HKY model, lognormal clock violations (σ = 0.5), biased sampling, cross-simulator checks (phastSim^33^), recombinant chimeras, dirty-data injection (high-N content, duplicates, misgrouping and date errors) and 1,500-tip scaling.

Clock-rate absolute relative error thresholds were 10% (strong signal) and 25% (moderate signal). Topological accuracy used unrooted normalized Robinson–Foulds distance (nRF) on shared tips^34^; tMRCA intervals used TreeTime 90% HPDs, with Clopper–Pearson 95% CIs for conditional coverage. For complete pairs passing EvoTrace’s pre-check tMRCA plausibility gate (root date ≤ earliest sample), LSD2 (v.2.4.4)^35^ analysed the same ML trees and dates; the paired median and 95% CI of log_2_(∣LSD2 relative error∣/∣EvoTrace relative error∣) were derived from 20,000 bootstraps (seed = 20261030).

Real-data benchmarks used six Nextstrain datasets^24^ (Zika, mumps, WNV, measles, dengue-1 and norovirus; downloaded 18 August 2026). Filtering to sequences in official Auspice trees (904–3,924 sequences), we ran three core replicates each (seeds = 1729, 2718 and 3141) and separately tested WNV untranslated region (UTR) masking. Metrics included nRF, official clade (≥3 tips) monophyly, patristic distance correlation (50,000 sampled tip pairs per run), root-to-tip slope and R^2^, and tMRCA (90% HPD).

### Three-way end-to-end comparison

Across the six frozen Nextstrain datasets, we benchmarked EvoTrace, Biomni^8^ and bare Kimi-K3—all powered by the Kimi-K3 backbone—using three random seeds (42, 12345 and 20260912; 32 threads, 24-h timeout per run; 54 runs in total).

Completion rates within 24 h and wall-clock times were evaluated across all 18 runs per method. Temporal accuracy was summarized over Zika, mumps, WNV, measles and dengue-1 (norovirus was restricted to completion and topology assessments due to weak temporal signal and genogroup pre-check differences). Cross-seed reproducibility was measured as the three-replicate root-date range; accuracy was quantified via root-date bias (run point estimate minus official regression intercept) and relative rate difference. We used exact two-sided Wilcoxon signed-rank tests to compare absolute root-date bias and wall-clock time (15 virus–seed pairs) and cross-seed ranges (5 dataset pairs), and Fisher’s exact test for completion rates.

### Evidence-chain reconstruction and controlled defect experiments

A run-level evidence chain is judged fully reconstructed when raw inputs and all declared artefacts (stage metrics, ML tree, time-tree manifest, dated time tree, branch lengths and clock summary) exist and their SHA-256 hashes match manifest reference hashes (report claims inherit their parent run’s status). In an isolated context, the independent auditor recomputes metrics and verifies plotting tables, arrays and graphical primitives, resolving and grading discrepancies by priority: raw artefacts > per-run metrics > summary tables > plotting tables > figures.

Controlled defect benchmarks used the 18 complete Nextstrain runs (blinded audit tolerance 1×10^−12^; two-sided 95% Wilson CIs for proportions). Structural chain tests included: (1) single-class defects (18 clean controls; 126 defective copies with artefact omissions, content drift or cross-case swaps); and (2) cumulative defects (tiers 0–4 sequentially stacking four drift/omission classes; 72 defective copies), defining the anomaly score as the fraction of eight required artefact classes that were missing or hash-unverified. Independent audit tests included: (1) overt defects (18 controls; 162 defective copies with tampered table, graphic or report values, or stale manifests), assessing detection, localization accuracy and clean acceptance; and (2) a numerical difficulty gradient (36 diagnostic and TreeTime R^2^ chains, each with one control and nine injected bias levels from 1×10^−5^ to 1×10^−2^; 360 trials). Maximizing Youden’s J index across dengue-1, measles and mumps (180 trials) yielded a threshold of 5.02×10^−5^, which was evaluated for detection and acceptance on the three held-out viruses.

### CCHFV phylogenetics, reassortment and ecological association analysis

CCHFV sequences from NCBI nuccore^36^ (28 August 2026) were filtered for sampling year, country and minimum length (S ≥ 1,500 bp, M ≥ 5,000 bp, L ≥ 11,000 bp). Quality control against reference strain IbAr 10200 required coverage ≥ 0.70, ambiguous bases ≤ 0.05, gaps ≤ 0.20 and coding completeness; exact deduplication (identity = 1.0) retained 432 (S), 399 (M) and 421 (L) sequences. Sequences were aligned with Nextclade (v.3.24.0)^37^, ML trees built via IQ-TREE (v.3.1.4)^38^ and ModelFinder^39^ (1,000 SH-aLRT^40^ and UFBoot^41^ replicates; midpoint-rooted), and time trees inferred using TreeTime (v.0.11.5; coalescent = opt, clock-filter-iqd = 4)^26^ and Augur refine^25^. Root-to-tip slope upper-tail significance used 100 tip-date permutations.

Average-linkage clustering on ML patristic distances maximized labelled-tip purity and minimized cluster count (6–30 clusters). Clusters were named sequentially: Europe 2 if containing Pentalofos-Greece-2015 or Malko Tarnovo-BG2012^42–44^; the majority label if ≥ 60% among labelled records; Europe 1 if unlabelled with ≥ 80% Turkish samples; or unclassified. Typing confidence (high, medium, low) reflected monophyly, label purity (1.0 or ≥ 0.8) and UFBoot (≥95%). Linking segments by normalized isolate name (retaining highest-confidence duplicates), isolates with ≥ 2 named segments were classified as candidate (discordant types), concordant (concordant_partial if with an unclassified segment) or uninformative. Candidate robustness was tested across 12 clustering-threshold perturbations.

Cross-border transmission was modelled via TreeTime mugration (parent–child country transitions with both confidences ≥ 0.70 counted as importations). A host-frequency sampling-bias correction factor (category count divided by Simpson’s effective category count, capped at 1–5) was applied. Topological robustness for major recipient countries was assessed by reconstructing ancestral states across 100 subsampled UFBoot trees (seed = 1729). Tick-genus association tests covered per-segment clusters and Europe 2 lineages with ≥ 10 pre-deduplication sequences and ≥ 5 tick records (23 tests). Fixing clusters, one-sided enrichment *P* values from 10,000 tick-label permutations (seed = 1729) were BH-adjusted^45^ and verified by exact hypergeometric tests. Paired analyses used complete pairs only: failed runs received no zero imputation, undelivered intervals exited conditional-coverage denominators, and unexecuted methods formed no valid pairs; all statistical tests were two-sided unless stated otherwise.

## Data availability

Source Data, frozen inputs, model outputs, run logs and audit materials supporting the findings of this study are archived by figure on GitHub (https://github.com/wuaipinglab/Auditable-AutoScience-EvoTrace). Material indices, file descriptions and access instructions are detailed in the repository documentation.

## Code availability

Code, configuration files and reproduction instructions for data processing, statistical analysis, figure generation and independent verification in this study are publicly available on GitHub (https://github.com/wuaipinglab/Auditable-AutoScience-EvoTrace). The repository provides figure-organized entry points, environment specifications and SHA-256 checksum manifests to reconstruct figure tables, recompute statistics and audit figures from frozen inputs and saved computational artefacts (full upstream re-execution additionally requires corresponding LLM API credentials, phylogenetic software environments and compute resources). The hosted multi-tenant web service layer of EvoTrace is maintained separately and accessible via https://evotrace.ai, with account access granted through the platform’s registration and approval process.

## Acknowledgements

This work was supported by the Prevention and Control of Emerging and Major Infectious Diseases–National Science and Technology Major Project (2025ZD01901800) to AW, the Major Project of Guangzhou National Laboratory (GZNL2024A01015) to AW, the National Natural Science Foundation of China (82550101) to AW and (32500551) to HZ, the CAMS Innovation Fund for Medical Sciences (CIFMS, 2025-I2M-XHXX-174) to HZ, and the CAMS Innovation Fund for Medical Sciences (CIFMS, 2026-I2M-SXZX7-02). The funders had no role in study design, data collection and analysis, decision to publish or preparation of the manuscript. Figure 1 (https://BioRender.com/eml5vb0), Fig. 5a (https://BioRender.com/cx3ncna) and Fig. 6a ( https://BioRender.com/1wpqqkw) were created in BioRender (Wu, A., 2026).

## Author notes

These authors contributed equally: Yun Ma, Hangyu Zhou.

These authors jointly supervised this work: Hangyu Zhou, Aiping Wu.

## Contributions

H.Z. and A.W. conceived and supervised the study. All authors designed the research. Y.M. and H.Z. collected the data and tested the models. Y.M. analysed the data, implemented the code and prepared the figures. H.Z. developed the web server. L.Q., Y.L. and J.Y. participated in the discussion. Y.M. wrote the manuscript, and H.Z. and A.W. revised it. All authors discussed the results and approved the final manuscript.

## Ethics declarations

## Competing interests

The authors declare no competing interests.

---

**Extended Data Fig. 1.**
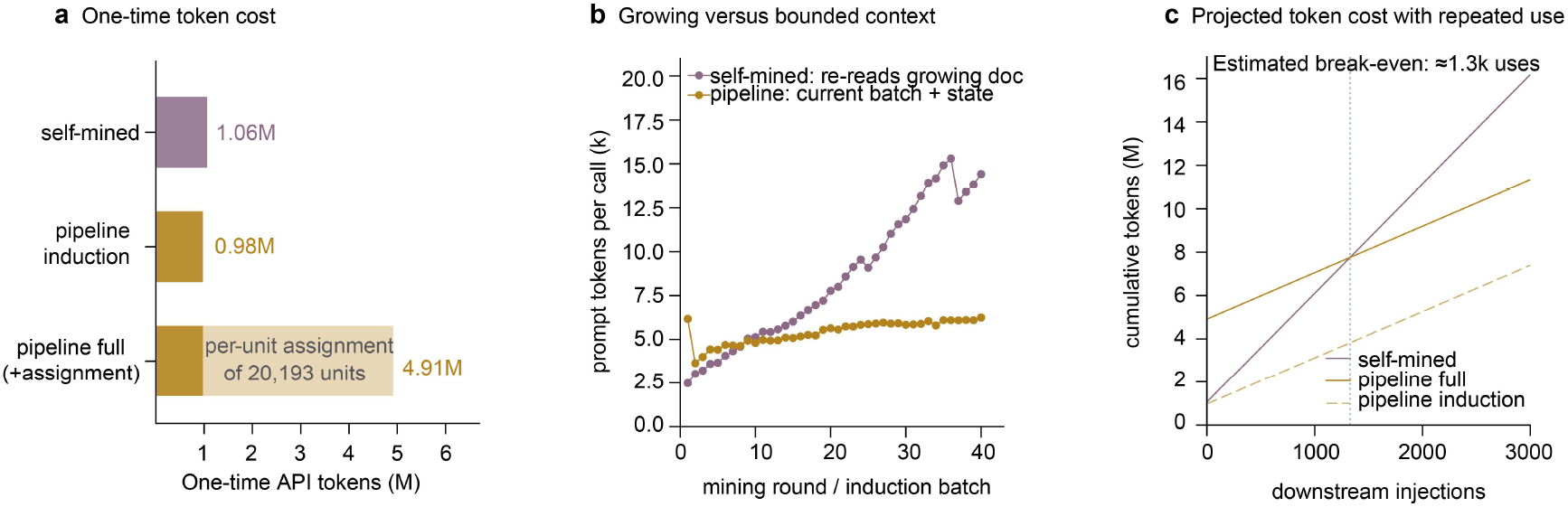
Token consumption for knowledge-base construction and downstream reuse. **a,** Total single-build token consumption for self-mined, pipeline-induction and full-pipeline modes. **b,** Per-call input token trajectories across 40 self-mined rounds and 40 induction batches. **c,** Extrapolated cumulative token consumption over repeated downstream calls (vertical dashed line, break-even point at ∼1,328 calls).

**Extended Data Fig. 2.**
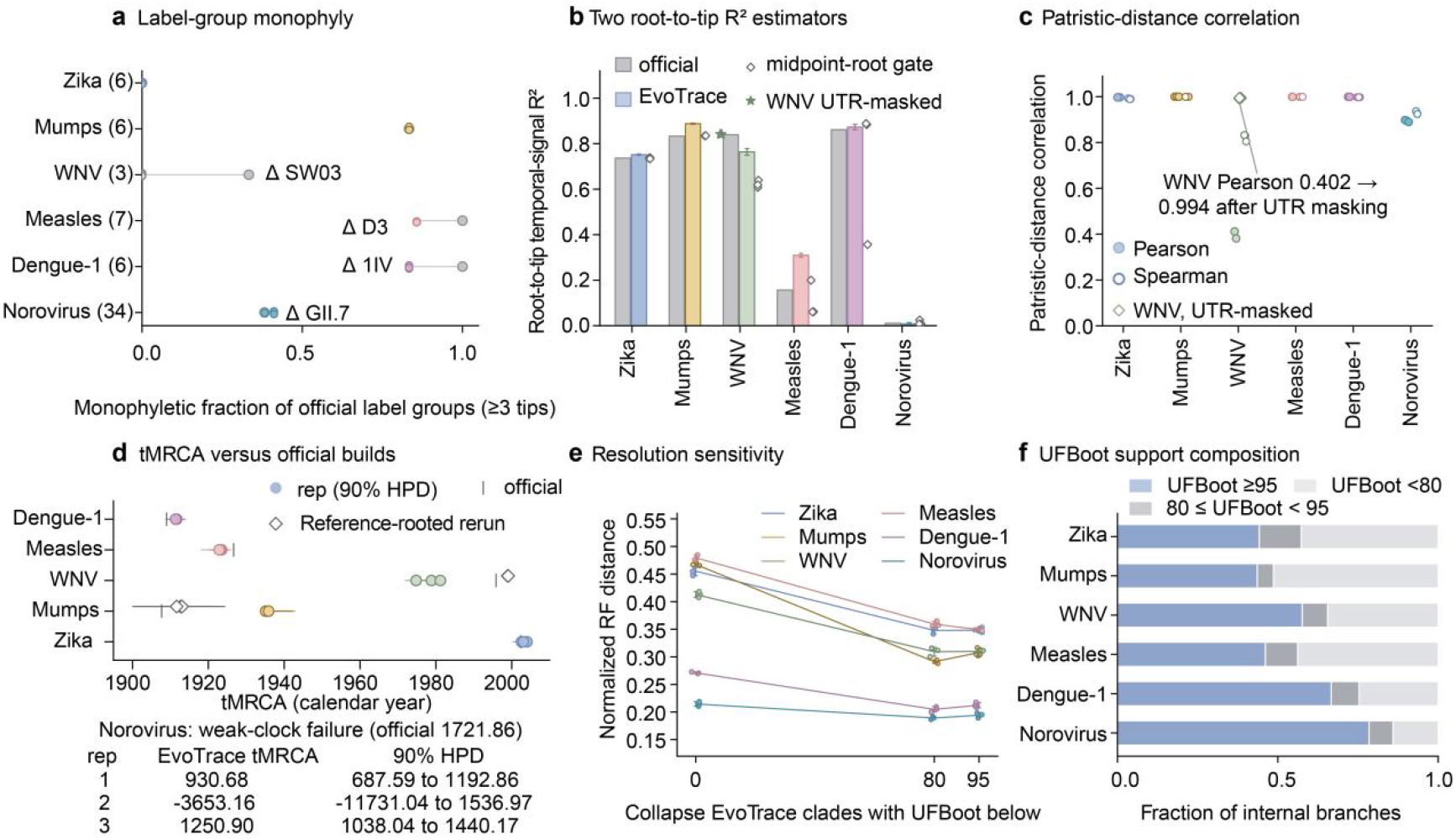
Consistency between EvoTrace and official Nextstrain benchmarks on empirical viral datasets (n = 3 independent runs per dataset). **a,** Monophyly retention proportions across 62 officially defined clades (triangles, non-monophyletic clades). **b,** Root-to-tip regression R^2^ (mean ± s.d.; diamonds, midpoint rooting; asterisks, recalculated after masking UTR noise). **c,** Pearson (solid) and Spearman (open) correlations of pairwise patristic distances between EvoTrace and official reference trees. **d,** tMRCA point estimates and 90% HPD intervals compared with official benchmarks (short bars; diamonds, recalculated with fixed identical roots). **e,** nRF topological distances (mean ± s.d.) across UFBoot branch-collapsing thresholds. **f,** Mean proportions of reconstructed internal nodes across three UFBoot support tiers.

**Extended Data Fig. 3.**
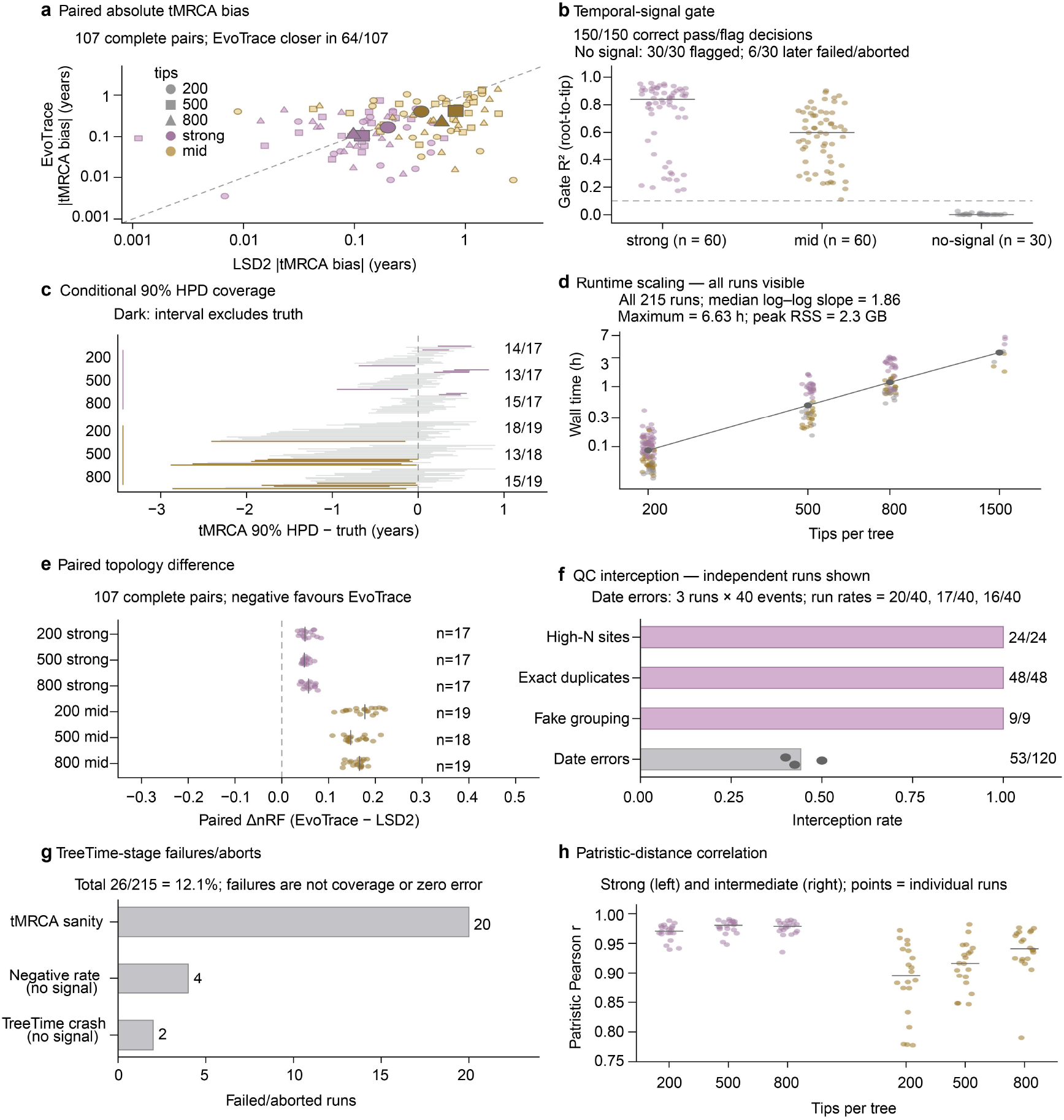
Ground-truth benchmarking, runtime scaling and gated quality control across 215 simulation runs. **a,** Paired tMRCA absolute errors between EvoTrace and LSD2 (n = 107; large circles, group medians; dashed line, y = x). **b,** Root-to-tip R^2^ distributions and temporal-signal gating (short bars, medians; dashed line, gate threshold; n = 60, 60 and 30 for strong, moderate and no signal). **c,** Ground-truth coverage calibration of tMRCA 90% HPD intervals across valid runs (n = 107; dark segments, uncalibrated/missed ground truth). **d,** Log–log power-law scaling of full-pipeline runtime versus sequence scale (tip count; n = 215; lines connect scale medians). **e,** Topological error differences (ΔnRF) between EvoTrace and LSD2 (negative values favour EvoTrace; n as in Fig. 3f). **f,** Interception rates for four pre-injected defect classes (high-N sites, duplicates, misgrouping and date errors). **g,** Attribution counts of runs intercepted by plausibility gates or aborted during TreeTime dating (26/215 runs). **h,** Pearson correlations of pairwise evolutionary distances between reconstructed and ground-truth trees (short bars, medians).

**Extended Data Fig. 4.**
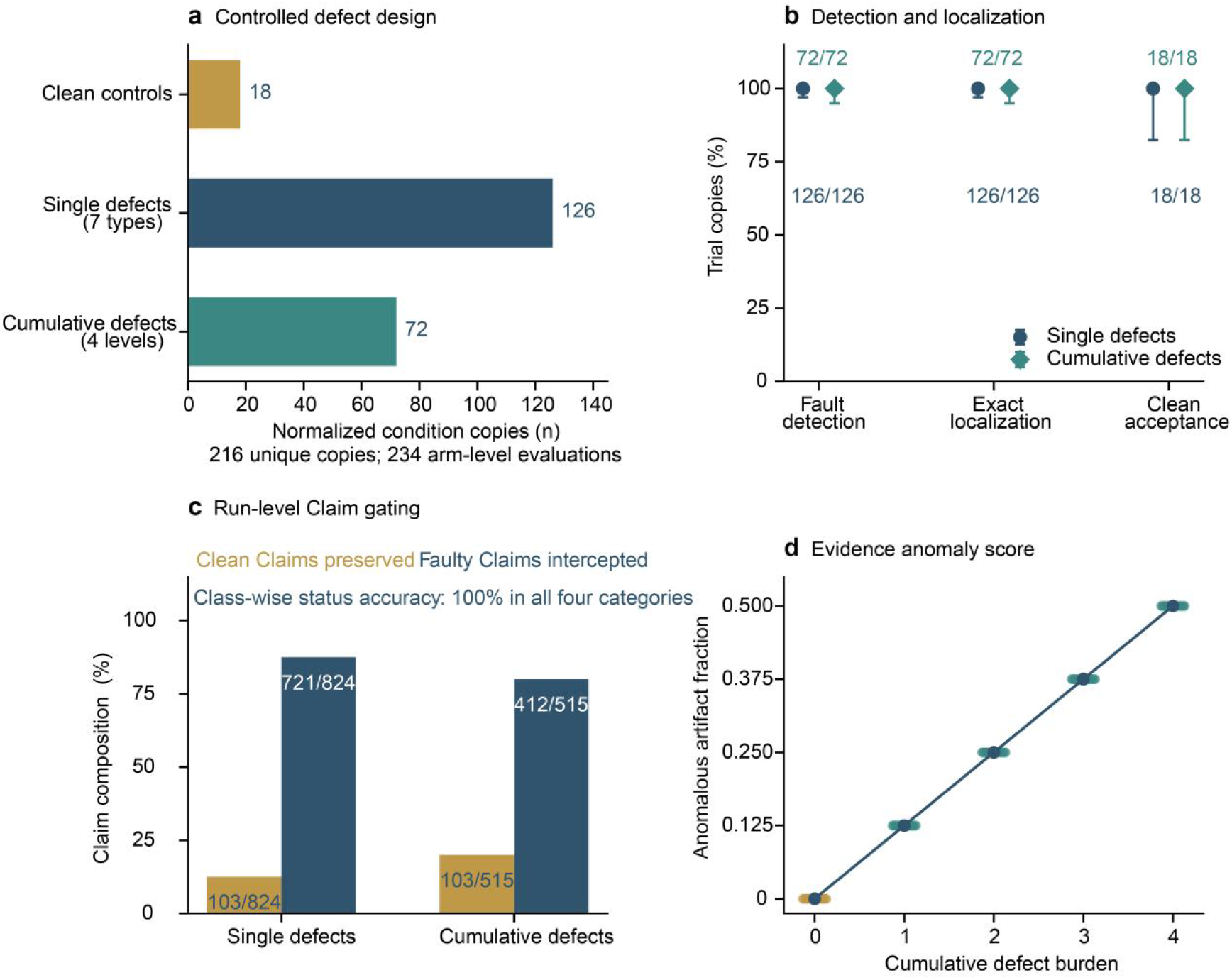
Verifiable evidence chains: anomaly detection, localization and claim gating under controlled defects. **a,** Two controlled experiments across 18 empirical analyses (234 paired-copy evaluations): single-defect arm (n = 144; 18 controls, 126 defect copies) and cumulative-defect arm (n = 90; 18 controls, 72 defect copies), totalling 216 unique copies after merging shared controls. **b,** Fault detection, pinpoint localization and clean acceptance under single-defect (dark blue, 126/126) and cumulative-defect (teal, 72/72) conditions, with 18/18 clean-control acceptance in both (error bars, 95% Wilson CIs). **c,** Bar heights show claim composition proportions. Clean and affected claims were 103/824 and 721/824 (single-defect) and 103/515 and 412/515 (cumulative-defect). Adjudication accuracy was 100% across all four claim categories (overall: 824/824 and 515/515; yellow, retained; blue, blocked; analysed independently per batch). **d,** Cumulative defect burden versus evidence anomaly score (proportion of eight mandatory artefacts missing or failing output checksum verification); burden levels 0–4 correspond to anomaly proportions of 0, 0.125, 0.250, 0.375 and 0.500 (n = 18 per level).

**Extended Data Fig. 5.**
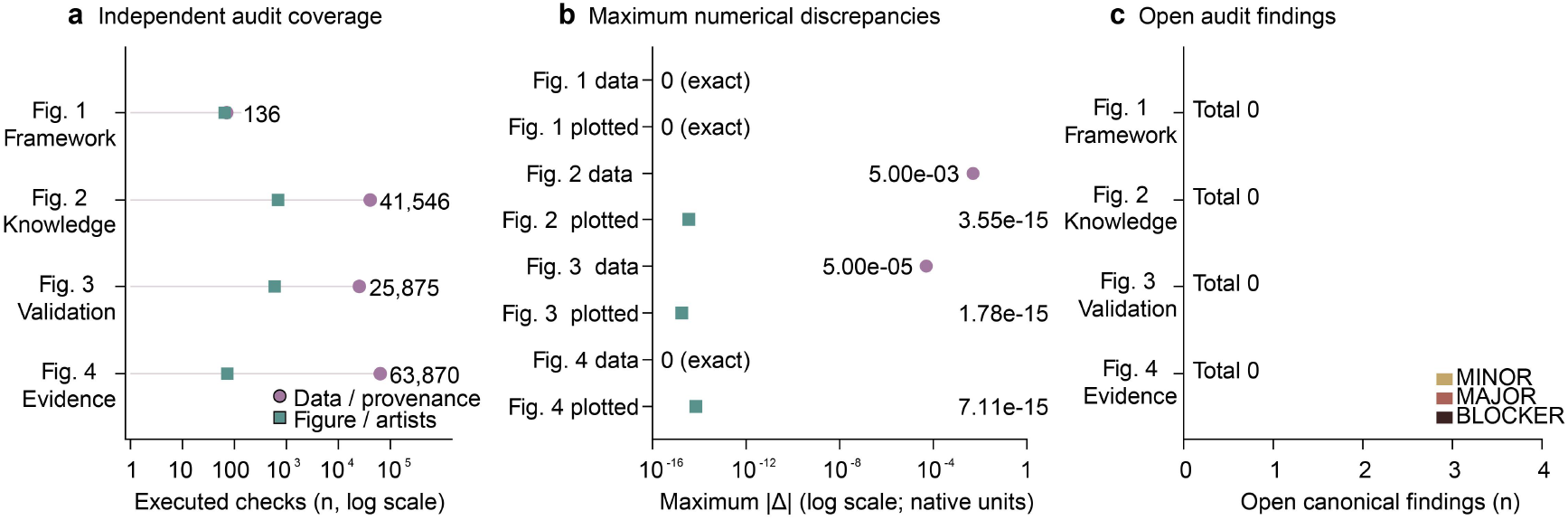
Independent re-audit performance after targeted remediation. **a,** Inspection-item counts from the independent re-audit of four remediated main-figure groups. **b,** Maximum absolute deviations across eight data-to-figure generation chains during re-audit. **c,** Unclosed issue counts across severity tiers upon independent re-audit (all reduced to zero).

**Extended Data Fig. 6.**
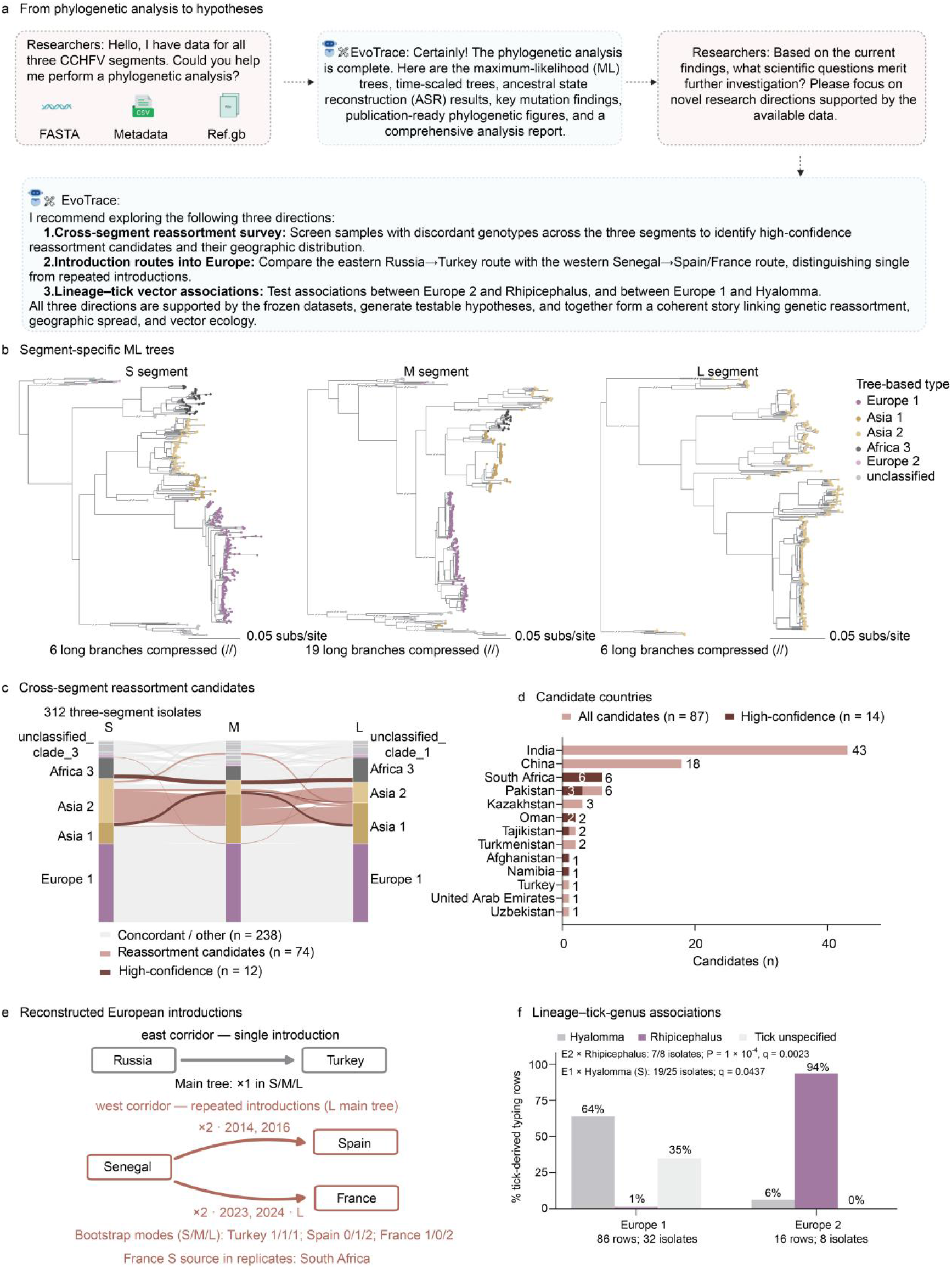
EvoTrace-enabled evolutionary analysis of CCHFV. **a,** Two-round researcher–EvoTrace interaction advancing from frozen-input phylogenetics to three linked hypotheses. **b,** Midpoint-rooted ML trees of S, M and L segments coloured by tree-based genotypes (branches > 0.08 substitutions per site compressed and marked //). **c,** S–M–L genotype combinations across 312 three-segment isolates: 74 reassortment candidates (12 high-confidence) and 238 non-candidates (199 concordant, 21 partially concordant, 18 indeterminate; unnamed cluster numbers are within-segment labels only). **d,** Country distribution of 87 reassortment candidates among 377 determinable multi-segment isolates (14 high-confidence: 12 three-segment, 2 two-segment; candidates require further validation). **e,** Dispersal pathways from mugration ancestral state reconstruction. Westward arrows show L-segment primary-tree transitions; modal counts are from 100 UFBoot replicates. Primary-tree versus replicate-mode differences: Spain S (1→0), M (2→1); France S (0→1; replicate source, South Africa). Dates are inferred; target tips (S/M/L) are 2/2/2 (Spain) and 5/2/10 (France). **f,** Tick-genus composition of pooled S, M and L records (≤3 records per isolate; light grey, unspecified genus, including 30/86 in Europe 1). Isolate-level associations used one-sided permutation tests (10,000 permutations) with unified BH correction across 23 tests: Europe 2–Rhipicephalus (7/8, *P*= 9.999×10−5, q = 0.0023); S-segment Europe 1–Hyalomma (19/25, q = 0.0437). Associations do not prove vector competence or rule out sampling/geographic confounding.

**Extended Data Fig. 7.**
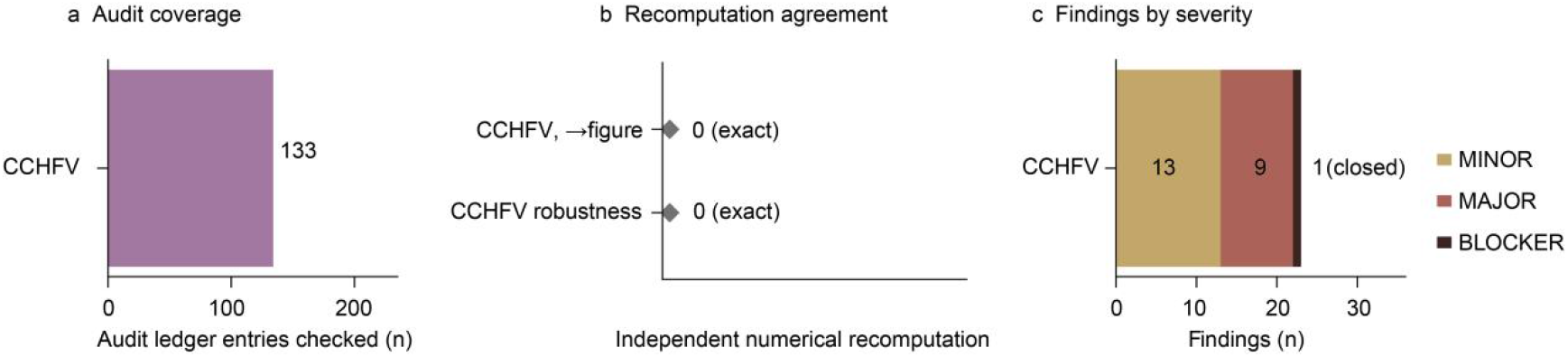
Independent audit of the CCHFV analytical evidence chain. **a,** Ten cross-version audits covering 133 un-deduplicated ledger entries (74+28+31; entries do not represent counts of independent scientific claims or confirmed biological truth). **b,** Independent numerical recomputation from source data to figure data and robustness analyses; both chains showed a maximum absolute difference of 0, explicitly marked as ‘0 (exact)’ without small-positive-value substitution. **c,** Cumulative audit findings: 13 MINOR, 9 MAJOR and 1 BLOCKER (excluding NOTE). The BLOCKER was confirmed closed upon re-audit (not implying closure of all other findings).

## Notes

https://evotrace.ai

https://github.com/wuaipinglab/Auditable-AutoScience-EvoTrace

### Competing Interest Statement

The authors have declared no competing interest.

## References

1. Wang, H. et al. Scientific discovery in the age of artificial intelligence. Nature 620, 47–60 (2023).

2. Guo, L. Y. et al. Toward autonomous science with agentic artificial intelligence. Cell 189, 6574–6588 (2026).

3. Boiko, D. A., MacKnight, R., Kline, B. & Gomes, G. Autonomous chemical research with large language models. Nature 624, 570–578 (2023).

4. Lu, C. et al. Towards end-to-end automation of AI research. Nature 651, 914–919 (2026).

5. Swanson, K., Wu, W., Bulaong, N. L., Pak, J. E. & Zou, J. The Virtual Lab of AI agents designs new SARS-CoV-2 nanobodies. Nature 646, 716–723 (2025).

6. Gottweis, J. et al. Accelerating scientific discovery with Co-Scientist. Nature 655, 487–496 (2026).

7. Ghareeb, A. E. et al. A multi-agent system for automating scientific discovery. Nature 655, 497–505 (2026).

8. Huang, K. et al. Autonomous biomedical research with an artificial intelligence agent. Science 393, eadz4351 (2026).

9. Miao, J., Davis, J. R., Zhang, Y., Pritchard, J. K. & Zou, J. Reimagining research papers as interactive and reliable AI agents. Nature 658, 1272–1280 (2026).

10. Kapoor, S. & Narayanan, A. Leakage and the reproducibility crisis in machine-learning-based science. Patterns 4, 100804 (2023).

11. Peng, R. D. Reproducible research in computational science. Science 334, 1226–1227 (2011).

12. Chen, Z. et al. ScienceAgentBench: toward rigorous assessment of language agents for data-driven scientific discovery. In Proc. 13th International Conference on Learning Representations https://openreview.net/forum?id=6z4YKr0GK6 (ICLR, 2025).

13. Starace, G. et al. PaperBench: evaluating AI’s ability to replicate AI research. In Proc. 42nd International Conference on Machine Learning Vol. 267, 56843–56873 (PMLR, 2025).

14. OpenAI. openai/math. GitHub https://github.com/openai/math (2026).

15. Gallegos, A. OpenAI withdraws three preprints a day after releasing 722 manuscripts on unsolved math problems. Retraction Watch https://retractionwatch.com/2026/10/08/openai-withdraws-preprints-722-manuscripts-unsolved-math-problems/ (2026).

16. Tao, T. Math 2.0. What’s new https://terrytao.wordpress.com/2026/10/10/math-2-0/ (2026).

17. Sandve, G. K., Nekrutenko, A., Taylor, J. & Hovig, E. Ten simple rules for reproducible computational research. PLoS Comput. Biol. 9, e1003285 (2013).

18. Stodden, V. et al. Enhancing reproducibility for computational methods. Science 354, 1240–1241 (2016).

19. Kassis, T., Agarwal, V., He, Y., Patel, D. & Brueckner, A. M. Scientific Agent Skills: a library of procedural knowledge for research agents. Preprint at 10.48550/arXiv.2609.00065 (2026).

20. Anthropic. Building agents: when to fold subagents into your main agent loop. Claude https://claude.com/resources/articles/building-agents-when-to-fold-subagents-into-your-main-agent-loop (2026).

21. Rambaut, A., Lam, T. T., Max Carvalho, L. & Pybus, O. G. Exploring the temporal structure of heterochronous sequences using TempEst (formerly Path-O-Gen). Virus Evol. 2, vew007 (2016).

22. Duchêne, S., Duchêne, D., Holmes, E. C. & Ho, S. Y. W. The performance of the date-randomization test in phylogenetic analyses of time-structured virus data. Mol. Biol. Evol. 32, 1895–1906 (2015).

23. Murray, G. G. R. et al. The effect of genetic structure on molecular dating and tests for temporal signal. Methods Ecol. Evol. 7, 80–89 (2016).

24. Hadfield, J. et al. Nextstrain: real-time tracking of pathogen evolution. Bioinformatics 34, 4121–4123 (2018).

25. Huddleston, J. et al. Augur: a bioinformatics toolkit for phylogenetic analyses of human pathogens. J. Open Source Softw. 6, 2906 (2021).

26. Sagulenko, P., Puller, V. & Neher, R. A. TreeTime: maximum-likelihood phylodynamic analysis. Virus Evol. 4, vex042 (2018).

27. Kosakovsky Pond, S. L. & Frost, S. D. W. Not so different after all: a comparison of methods for detecting amino acid sites under selection. Mol. Biol. Evol. 22, 1208–1222 (2005).

28. Baele, G. et al. BEAST X for Bayesian phylogenetic, phylogeographic and phylodynamic inference. Nat. Methods 22, 1653–1656 (2025).

29. Gill, M. S. et al. Improving Bayesian population dynamics inference: a coalescent-based model for multiple loci. Mol. Biol. Evol. 30, 713–724 (2013).

30. Bouckaert, R. et al. BEAST 2.5: an advanced software platform for Bayesian evolutionary analysis. PLoS Comput. Biol. 15, e1006650 (2019).

31. Müller, N. F., Rasmussen, D. A. & Stadler, T. MASCOT: parameter and state inference under the marginal structured coalescent approximation. Bioinformatics 34, 3843–3848 (2018).

32. Ly-Trong, N., Naser-Khdour, S., Lanfear, R. & Minh, B. Q. AliSim: a fast and versatile phylogenetic sequence simulator for the genomic era. Mol. Biol. Evol. 39, msac092 (2022).

33. De Maio, N. et al. phastSim: efficient simulation of sequence evolution for pandemic-scale datasets. PLoS Comput. Biol. 18, e1010056 (2022).

34. Robinson, D. F. & Foulds, L. R. Comparison of phylogenetic trees. Math. Biosci. 53, 131–147 (1981).

35. To, T.-H., Jung, M., Lycett, S. & Gascuel, O. Fast dating using least-squares criteria and algorithms. Syst. Biol. 65, 82–97 (2016).

36. Sayers, E. W. et al. Database resources of the National Center for Biotechnology Information. Nucleic Acids Res. 50, D20–D26 (2022).

37. Aksamentov, I., Roemer, C., Hodcroft, E. B. & Neher, R. A. Nextclade: clade assignment, mutation calling and quality control for viral genomes. J. Open Source Softw. 6, 3773 (2021).

38. Minh, B. Q. et al. IQ-TREE 2: new models and efficient methods for phylogenetic inference in the genomic era. Mol. Biol. Evol. 37, 1530–1534 (2020).

39. Kalyaanamoorthy, S., Minh, B. Q., Wong, T. K. F., von Haeseler, A. & Jermiin, L. S. ModelFinder: fast model selection for accurate phylogenetic estimates. Nat. Methods 14, 587–589 (2017).

40. Guindon, S. et al. New algorithms and methods to estimate maximum-likelihood phylogenies: assessing the performance of PhyML 3.0. Syst. Biol. 59, 307–321 (2010).

41. Hoang, D. T., Chernomor, O., von Haeseler, A., Minh, B. Q. & Vinh, L. S. UFBoot2: improving the ultrafast bootstrap approximation. Mol. Biol. Evol. 35, 518–522 (2018).

42. Papa, A. et al. Isolation and whole-genome sequencing of a Crimean-Congo hemorrhagic fever virus strain, Greece. Ticks Tick Borne Dis. 9, 788–791 (2018).

43. Panayotova, E., Papa, A., Trifonova, I. & Christova, I. Crimean-Congo hemorrhagic fever virus lineages Europe 1 and Europe 2 in Bulgarian ticks. Ticks Tick Borne Dis. 7, 1024–1028 (2016).

44. Hua, B. L. et al. A single mutation in Crimean-Congo hemorrhagic fever virus discovered in ticks impairs infectivity in human cells. eLife 9, e50999 (2020).

45. Benjamini, Y. & Hochberg, Y. Controlling the false discovery rate: a practical and powerful approach to multiple testing. J. R. Stat. Soc. B 57, 289–300 (1995).

